# Binding of inhibitory checkpoints to CD18 in *cis* hinders anti-cancer immune responses

**DOI:** 10.1101/2025.09.10.675342

**Authors:** Zhenghai Tang, Ming-Chao Zhong, Jin Qian, Lok San Wong, Jiaxin Li, Dominique Davidson, André Veillette

**Author notes:** Correspondence (Z.T.) and (A.V.).

## Abstract

SIRPα is an inhibitory receptor on macrophages that limits phagocytosis and anti-tumor activity of macrophages by “*trans*” interacting with CD47 on tumor cells. Herein, we found that a large component of SIRPα’s inhibitory function occurred independently of CD47-binding and phosphatase signaling. This function resulted from a direct interaction between SIRPα and CD18 (β2 integrin) in “*cis*” at the surface of macrophages, involving SIRPα amino acids distinct from those implicated in the SIRPα-CD47 interaction. The *cis* interaction prevented activation of CD18, which is necessary for phagocytosis. Combined blockade of SIRPα-CD18 and SIRPα-CD47 was essential for maximizing phagocytosis and suppression of tumor growth *in vivo*. Similar *cis* interactions between CD18 and other inhibitory checkpoints, including PD-1, were also observed. Thus, in addition to mediating effects when engaged by ligands in *trans*, inhibitory checkpoints suppress immune cell activation through a mechanism targeting CD18 in *cis*. This dual mode of action should be considered when developing blockers of inhibitory checkpoints for immunotherapy.

**One-Sentence Summary:** In addition to being engaged in “*trans*” by ligands on tumor cells, inhibitory receptors, such as SIRPα and PD-1, hinder anti-cancer immune responses by “*cis*” interacting with β2 integrin CD18.

---

Engagement of inhibitory immune checkpoints in “*trans*” by ligands on cancer cells or in the tumor microenvironment can severely compromise anti-cancer immunity, favoring tumor progression (*1–3*). This observation led to the development of drugs, such as monoclonal antibodies (mAbs) against the PD-1 and CTLA-4 receptors, blocking these interactions to promote anti-tumor immunity and have shown considerable clinical efficacy (*4, 5*). However, many patients fail to respond or experience relapse. To improve efficacy, additional efforts have focused on other immune checkpoints expressed on T cells or other immune cells, including macrophages.

SIRPα is the most extensively studied inhibitory receptor found on macrophages, which enable destruction of cancer cells by phagocytosis (*2, 6–13*). Upon engagement by its ligand CD47, often overexpressed on cancer cells, SIRPα transmits inhibitory signals through intracellular phosphatases that limit phagocytosis (*6, 7, 12–15*). Unfortunately, disruption of the SIRPα-CD47 interaction using SIRPα or CD47 blocking agents has produced variable results in early clinical trials (*9, 10*). One possibility is that the strategy for targeting SIRPα is not optimal, perhaps due to an incomplete understanding of the mechanisms of action of SIRPα. The same caveat may apply to other inhibitory checkpoints in other cell types.

## SIRPα suppresses phagocytosis independently of CD47 and phosphatase signaling

Current models indicate that, when SIRPα is engaged in *trans* by CD47 on tumor cells, the intracellular immunoreceptor tyrosine-based inhibitory motifs (ITIMs) of SIRPα recruit the protein tyrosine phosphatases SHP-1 and SHP-2, which suppress activating signals emanating from pro-phagocytic receptors such as Fc receptors (FcRs) and SLAMF7 (**Fig. 1 A**). To investigate whether SIRPα’s inhibitory function was solely dependent on CD47-binding and phosphatase signaling, various SIRPα mutants were introduced via retroviral infection into bone marrow-derived macrophages (BMDMs) from SIRPα-deficient (knock-out; KO) mice (**Fig. 1, B and C and fig. S1A**). These mutants included SIRPα^T96V^, which involves a residue critical for binding to CD47 (L66 in the human SIRPα V1 sequence from reference (*16*)); a variant carrying tyrosine-to-phenylalanine mutations within the four ITIMs (SIRPα^FFFF^); and a mutant lacking most of the cytoplasmic domain (SIRPα^ΔIC^)—alone or in combination. All variants were expressed at levels comparable to wild-type (WT) SIRPα, and those with the T96V mutation failed to bind a CD47-Fc fusion protein, compared to WT SIRPα (**Fig. 1D**).

**Fig. 1.**
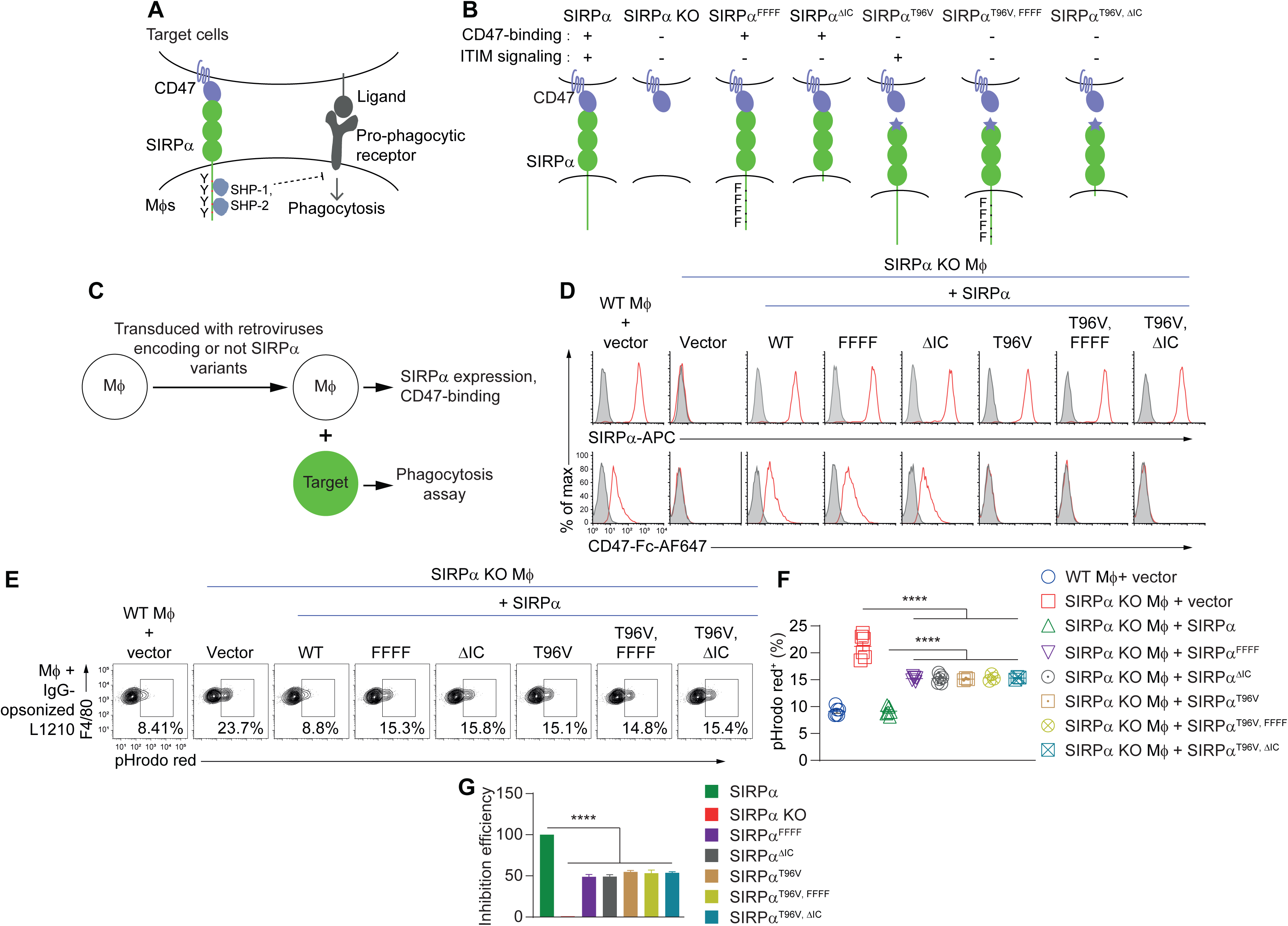
SIRPα suppresses phagocytosis independently of CD47 and phosphatase signaling. (**A**) Existing model by which SIRPα suppresses phagocytosis by interacting in *trans* with CD47 on target cells. See text for details. The 3 Ig-like domains of SIRPα (1 IgV and 2 IgCs) and the single Ig-V domain of CD47 are shown as ellipses. Mβs, macrophages. (**B**) Depiction of SIRPα variants and their functional characteristics. SIRPα^FFFF^ contained substitution of tyrosine (Y)-to-phenylalanine (F) substitution at Y436, 460, 477, and 501; SIRPα^ΔIC^ lacked most of the cytoplasmic domain of SIRPα, ending with arginine 401; SIRPα^T96V^ carried a threonine (T)-to-valine (V) mutation at position 96 (shown by lavender star), which abolishes CD47-binding; SIRPα^T96V,FFFF^ had the T96V and FFFF mutations; SIRPα^T96V,ΔIC^ had the T96V and the ΔIC mutations. KO, knock-out. ITIM, immunoreceptor tyrosine-based inhibitory motif. (**C** to **G**) SIRPα variants or empty vector were expressed in SIRPα KO BMDMs and tested. Wild-type (WT) BMDMs were used as control. (**C**) Schematic representation of assays performed. Fc, fragment crystallizable. (**D**) Flow cytometry analyses of SIRPα expression and CD47-binding. APC, allophycocyanin. AF647, Alexa fluor 647. (**E** and **F**) Representative (**E**) and compiled data (**F**) of pHrodo-based phagocytosis assays using L1210 derivatives expressing Tac and opsonized with Tac monoclonal antibody (mAb) 7G7, as targets. Positive cells with percentages are boxed. **G**, Efficiency of phagocytosis inhibition in SIRPα KO BMDMs expressing or not the indicated SIRPα variants was calculated using the values in (**F**). SIRPα KO expressing WT SIRPα or empty vector displayed 100% and 0% inhibition efficiency, respectively. All data are means ± s.e.m., \*\*\*\**p* < 0.0001. Results in (**D** and **E**) are representative of 6 independent experiments, except for SIRPα^T96V^, SIRPα^T96V, FFFF^ and SIRPα^T96V, ΔIC^ that are representative of 3 experiments. Results in (**F** and **G**) are pooled from a total of 6 mice studied in 6 independent experiments, except for SIRPα^T96V^, SIRPα^T96V, FFFF^ and SIRPα^T96V, ΔIC^ that involved 3 mice in 3 experiments. Each symbol in (**F**) represents one mouse.

In a pHrodo-based phagocytosis assay with IgG-opsonized mouse leukemia cells L1210 as targets, SIRPα KO BMDMs exhibited increased phagocytosis compared to WT BMDMs, as previously described (*17*) **(Fig. 1, E to G, and fig. S1, B and C**). This increase was fully reversed by re-expression of WT SIRPα, but all mutants—SIRPα^FFFF^, SIRPα^ΔIC^, SIRPα^T96V^, SIRPα^T96V, FFFF^, and SIRPα^T96V, ΔIC^—only partially suppressed phagocytosis (by ∼50%) (**Fig. 1, E to G**). Similar results were observed in a microscopy-based phagocytosis assay, using IgG-opsonized L1210, EL-4 (lymphoma), or P815 (mastocytoma) cells as targets (**fig. S1D**).

Therefore, a substantial component of SIRPα’s capacity to suppress phagocytosis was independent of CD47-binding and phosphatase signaling.

## SIRPα binds to integrin CD18 in *cis*

SIRPα may mediate additional inhibitory effects by influencing one or more partner(s) at the plasma membrane. To address this possibility, SIRPα was immunoprecipitated from WT BMDMs, and potential SIRPα-associated proteins were identified by mass spectrometry (**Fig. 2, A and B**). Several plasma membrane proteins, including CD45, CD18 (β2 integrin), CD11b (α_M_ integrin), ATPase subunit alpha-1 (ATP1A1) and EGF-like module-containing mucin-like hormone receptor-like 1 (EMR1) were immunoprecipitated from cells expressing SIRPα, but not from cells lacking SIRPα (**Fig. 2B and fig. S2A**). Among these proteins, CD45 is a transmembrane protein tyrosine phosphatase with a reported ability to regulate macrophage functions (*18*). However, little or no difference in phagocytosis of IgG-opsonized L1210 cells was observed in WT or SIRPα KO BMDMs lacking CD45, compared to their CD45-expressing counterparts (**fig. S2, B and C**).

**Fig. 2.**
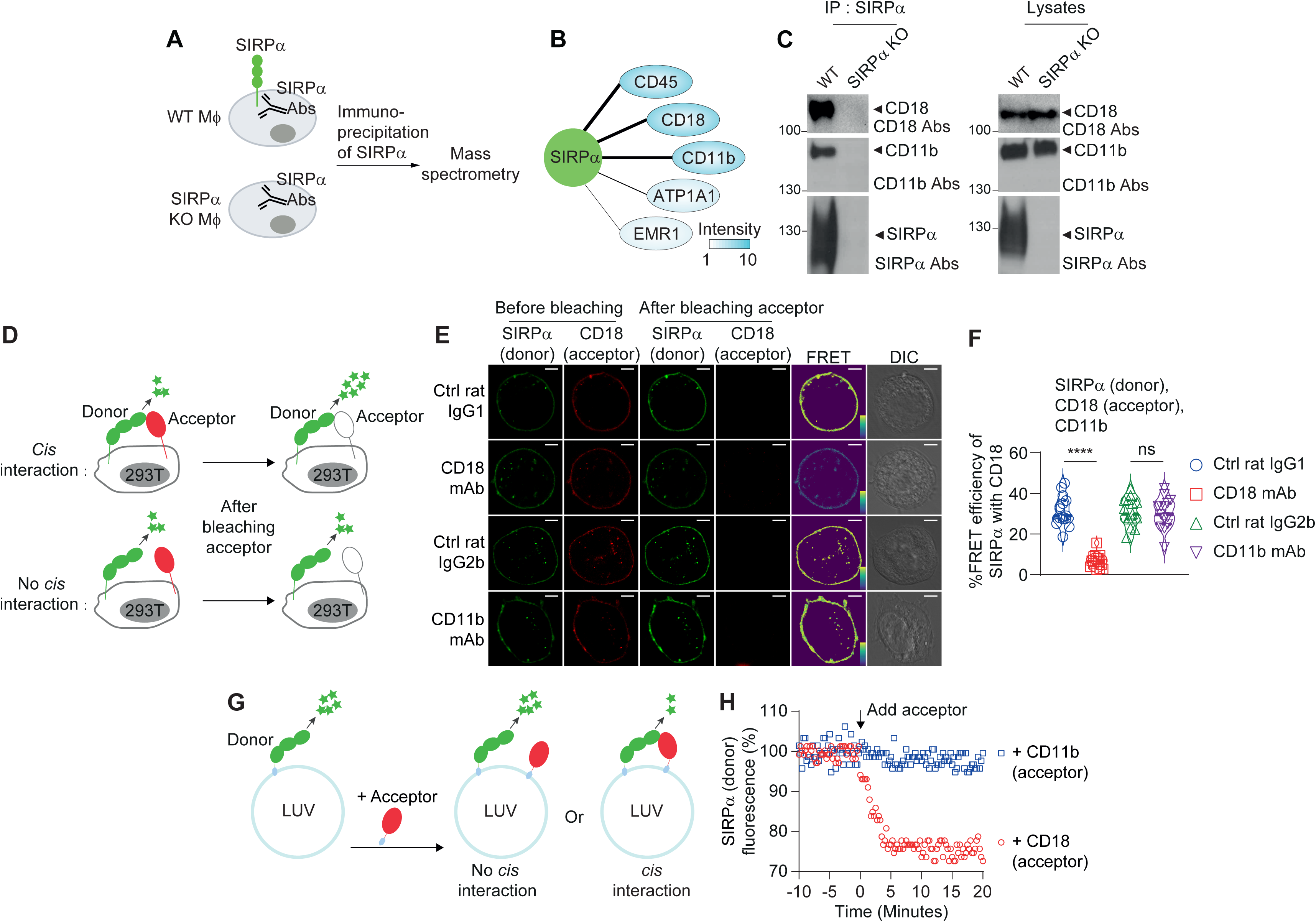
SIRPα binds to integrin CD18 in *cis* in macrophages. (**A** and **B**) Immunoprecipitation followed by mass spectrometry of SIRPα-associated proteins. (**A**) Schematic representation of assay. (**B**) Plasma membrane-associated proteins found in SIRPα immunoprecipitates from WT BMDMs, but not from SIRPα KO BMDMs. **c**, Co-immunoprecipitation assay of SIRPα, CD18 and CD11b in WT and SIRPα KO BMDMs. IP, immunoprecipitation. Abs, antibodies. (**D** to **F**) FRET assays. (**D**) Schematic representation of FRET assay in HEK293T cells. (**E** and **F**) Representative confocal microscopy images (**E**) and compiled data (**F**) of FRET assays with donor-labeled SIRPα, acceptor-labeled CD18 and unlabeled CD11b in the presence of control (Ctrl) IgG, CD18 mAb GAME-46 or CD11b mAb 5C6. Yellow to purple spectrum denotes strong to weak FRET. DIC, differential interference contrast. Scale bars, 5 μm. (**G** and **H**) LUV-FRET assay. (**G**) Schematic representation of LUV-FRET assay. (**H**), Time-course of donor-labeled SIRPα fluorescence intensity after addition of acceptor-labeled CD18 or CD11b, monitored with a real-time plate reader. All data are means ± s.e.m. ns, not significant, \*\*\*\**p* < 0.0001. Results in (**C**, **E** and **H**) are representative of 3 independent experiments. Results in (**B** and **F**) are pooled from a total of 3 independent experiments. Each symbol in (**F**) represents one cell.

CD18 is an integrin crucially involved in immune cell activation, adhesion, and migration (*19–21*). In macrophages, CD18 and CD11b form a complex known as Mac-1 or αMβ2. CD18 is required for stable plasma membrane expression of CD11b, and mediates Mac-1 signaling by “inside-out” and by “outside-in” mechanisms (*13, 22–24*). CD11b has up to 40 potential ligands, including intercellular adhesion molecule (ICAM)-1 and ICAM-2, which trigger the function of Mac-1. Consistent with the critical role of Mac-1 in phagocytosis, blockade of CD18 or CD11b with mAbs, or genetic loss of CD18, greatly reduced phagocytosis of IgG-opsonized L1210 cells by SIRPα KO BMDMs, compared to controls (**fig. S2, D to F**).

To test if Mac-1 was the target for CD47-and phosphatase signaling-independent inhibition by SIRPα, we assessed if SIRPα and Mac-1 were part of a *bona fide* complex at the plasma membrane, using several approaches. Immunoprecipitation experiments with BMDMs followed by immunoblots confirmed the co-immunoprecipitation of SIRPα with CD18 or CD11b (**Fig. 2C**). We also studied fluorescence resonance energy transfer (FRET), which can be detected when two molecules are positioned within 10 nm of each other and is revealed by an augmentation of “donor” fluorescence after photobleaching of the “acceptor” (*25*) (**Fig. 2D**). Studies using donor-labeled SIRPα and acceptor-labeled CD18, plus an unlabeled version of CD11b for stable expression of CD18, showed that photobleaching of CD18 increased the fluorescence of SIRPα, compared to the non-photobleached condition, indicating energy transfer from SIRPα to CD18 (**Fig. 2, E and F**). Similar results were obtained when cells expressed a donor-labeled SIRPα and an acceptor-labeled CD11b, plus an unlabeled CD18 (**fig. S3, A and B**). FRET between SIRPα and CD18 was also seen when SIRPα was co-expressed with CD18 and CD11a to form LFA-1 (also known as αLβ2), which is required for T cell and NK cell activation (*26*) (**fig. S3, C and D)**. FRET between SIRPα and CD18 was reduced by CD18 mAbs, but not by CD11b mAbs, compared to control (Ctrl) IgG (**Fig. 2, E and F**). However, CD11b mAbs, but not CD18 mAb, prevented FRET between CD11b and SLAMF7, supporting our previous finding that SLAMF7 mediates its pro-phagocytic function by interacting with CD11b (*13*) (**fig. S3, E to G**).

To study whether the physical proximity of SIRPα and CD18 was due to a direct interaction, FRET assays were performed using cell-free large unilamellar vesicles (LUVs) (*17*), which were reconstituted with donor-labeled SIRPα and acceptor-labeled CD18 or CD11b (**Fig. 2G**). Addition of CD18, but not of CD11b, quenched the fluorescence of SIRPα, indicating that SIRPα bound directly to CD18, but not to CD11b (**Fig. 2H**).

Hence, SIRPα directly interacted in *cis* with CD18, but not with CD11b (**fig. S3G**). In turn, CD11b interacted with the pro-phagocytic receptor SLAMF7.

## SIRPα utilizes different mechanisms for binding to CD18 and CD47

In order to determine the requirements for binding of SIRPα to CD18, we investigated whether CD18 interacted with SIRPβ1a, a SIRP family member that shares structural and sequence similarities with SIRPα, yet does not interact with CD47 (*6, 27*) (**Fig. 3A, and fig. S4A**). In FRET assays, there was minimal energy transfer between SIRPβ1a and CD18, compared to SIRPα and CD18, suggesting little to no *cis* interaction of SIRPβ1a with CD18 (**Fig. 3B, and fig. S4B**).

**Fig. 3.**
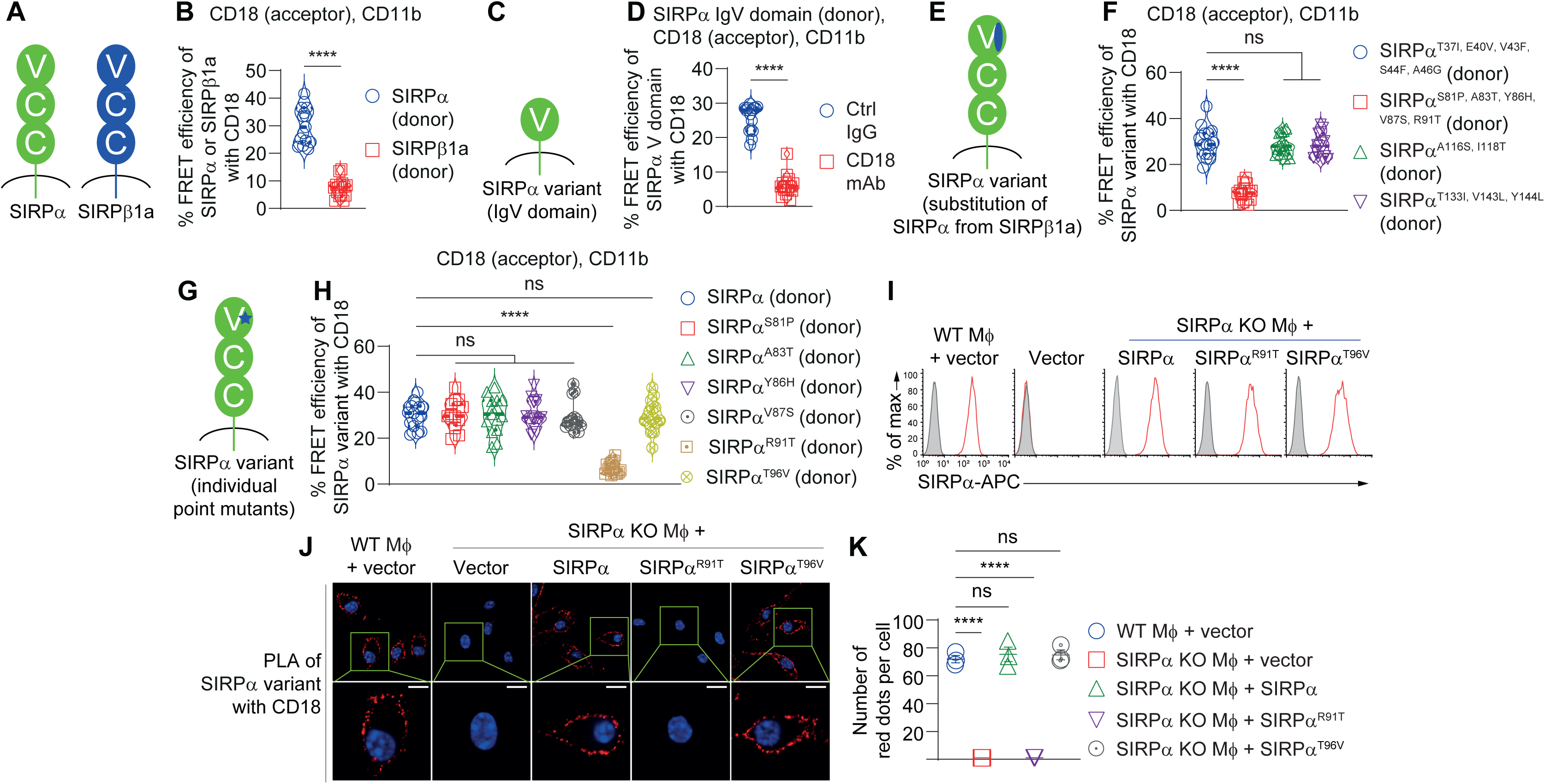
SIRPα utilizes different mechanisms for binding to CD18 and CD47. (**A** and **B**) FRET assays with SIRPα and SIRPβ1a. (**A**) A schematic representation of SIRPα and SIRPβ1a, with their 1 IgV domain and 2 IgC domains, is depicted. (**B**) Compiled data of 3 independent experiments using donor-labeled SIRPα or SIRPβ1a, acceptor-labeled CD18 and unlabeled CD11b, as done for Fig. 2, D to F. (**C** and **D**) FRET assays using SIRPα IgV domain. (**C**) A schematic representation of a SIRPα variant having only the IgV domain is shown. (**D**) Compiled data of 3 independent experiments using donor-labeled SIRPα IgV, acceptor-labeled CD18 and unlabeled CD11b, as done for Fig. 2, D to F. (**E**-**H**) FRET assays using SIRPα variants carrying non-conserved residues from SIRPβ1a. (**E** and **G**) Schematic representations of SIRPα variants. (**F** and **H**) Compiled data from 3 independent experiments using donor-labeled SIRPα variants, acceptor-labeled CD18 and unlabeled CD11b, as done for Fig. 2, D to F. (**I** to **K**) Proximity ligation assay (PLA) of SIRPα and CD18 in BMDMs expressing or not the indicated SIRPα variants. (I) Flow cytometry analyses of SIRPα expression. (**J** and **K**) Representative confocal microscopy images (**J**) and compiled data from 3 independent experiments (**K**) of PLA for SIRPα and CD18. Scale bar, 10 μm. All data are means ± s.e.m. ns, not significant, \*\*\*\**p* < 0.0001. Results in (**I** and **J**) are representative of 3 independent experiments. Results in (**B**, **D**, **F**, **H** and **K**) are pooled from 3 independent experiments. Each symbol in (**B**, **D**, **F**, **H** and **K**) represents one cell or mouse.

As CD18 interacted with a variant of SIRPα that contained only the IgV domain, which is known to bind CD47 (*8*) (**Fig. 3, C and D, and fig. S4C**), we substituted non-conserved amino acids in the SIRPα IgV domain by the corresponding residues in SIRPβ1a and assessed the SIRPα-CD18 interaction. SIRPα^S81P, A83T, Y86H, V87S, R91T^ exhibited a strong reduction of energy transfer with CD18, while SIRPαT37I, E40V, V43F, S44F, A46G, SIRPα^A116S, I118T^ and SIRPα^T133I, V143L, Y144L^ did not, when compared to WT SIRPα (**Fig. 3, E and F, and fig. S4D**). By creating individual substitutions, we found that SIRPα^R91T^, unlike SIRPα^S81P^, SIRPα^A83T^, SIRPα^Y86H^, or SIRPα^V87S^, and unlike SIRPα^T96V^, failed to interact with CD18 (**Fig. 3, G and H, and fig. S4, E and F**). The SIRPα^R91T^ mutation also inhibited energy transfer in the LUV-FRET assay (**fig. S4G**). The role of R91 in the SIRPα-CD18 interaction was also ascertained in BMDMs, using a proximity ligation assay (PLA). This technique results in the formation of fluorescent dots when two molecules are within 40 nm of one another (*17*). Fluorescent dots were observed in WT BMDMs, or in SIRPα KO BMDMs reconstituted with WT or SIRPα^T96V^, but not in SIRPα KO BMDMs expressing SIRPα^R91T^ or empty vector (**Fig. 3, J and K**). Unlike T96V, the R91T mutation did not impact binding to CD47-Fc (**fig. S5, A to C**, **and fig. S6A**).

Since T96, which is part of the CD47-binding site, and R91 are only five amino acids apart, we explored the possibility of competition between CD18 and CD47 for binding to SIRPα, through various methods. In a displacement assay, the SIRPα-CD18 complex identified by PLA was not disrupted by soluble CD47-His fusion proteins (**fig. S5, D and E**). Furthermore, co-expression of Mac-1 in SIRPα-expressing HEK293T cells did not affect binding of CD47-Fc, compared to cells not co-expressing Mac-1 (**fig. S5, F to I**). Analogous results were observed when CD47-Fc was used to stain WT or CD18 KO BMDMs (**fig. S5, J to L**).

Therefore, distinct SIRPα residues were involved in the interactions with CD18 and CD47.

## SIRPα-CD18 suppresses Mac-1 activation needed for phagocytosis

Next, we investigated whether the interaction of SIRPα with CD18 was restricting phagocytosis, a process dependent on Mac-1 activation. To this end, SIRPα KO BMDMs were reconstituted with WT SIRPα and SIRPα variants that lacked the *cis* interaction (SIRPα^R91T^), the *trans* interaction (SIRPα^T96V^), or the ability to transmit phosphatase signaling (SIRPα^FFFF^), individually or in combination (**Fig. 4a**). All SIRPα variants were expressed at levels comparable to WT SIRPα (**fig. S6A**). Their capacity to bind CD47 was also similar except for BMDMs carrying the SIRPα^T96V^ variants, which lost the ability to bind CD47-Fc, as expected (**fig. S6A**). In phagocytosis assays, SIRPα^R91T^ exhibited a reduced ability to suppress phagocytosis of IgG-opsonized L1210, EL-4, and P815 cells compared to WT SIRPα, (**Fig. 4, B and C, and fig. S6, B to E**). The R91T mutation also compromised further the inhibitory capacities of SIRPα^T96V^ and SIRPα^FFFF^, enhancing phagocytosis to levels seen with SIRPα deficiency (**Fig. 4, B and C, and fig. S6, B to E**).

**Fig. 4.**
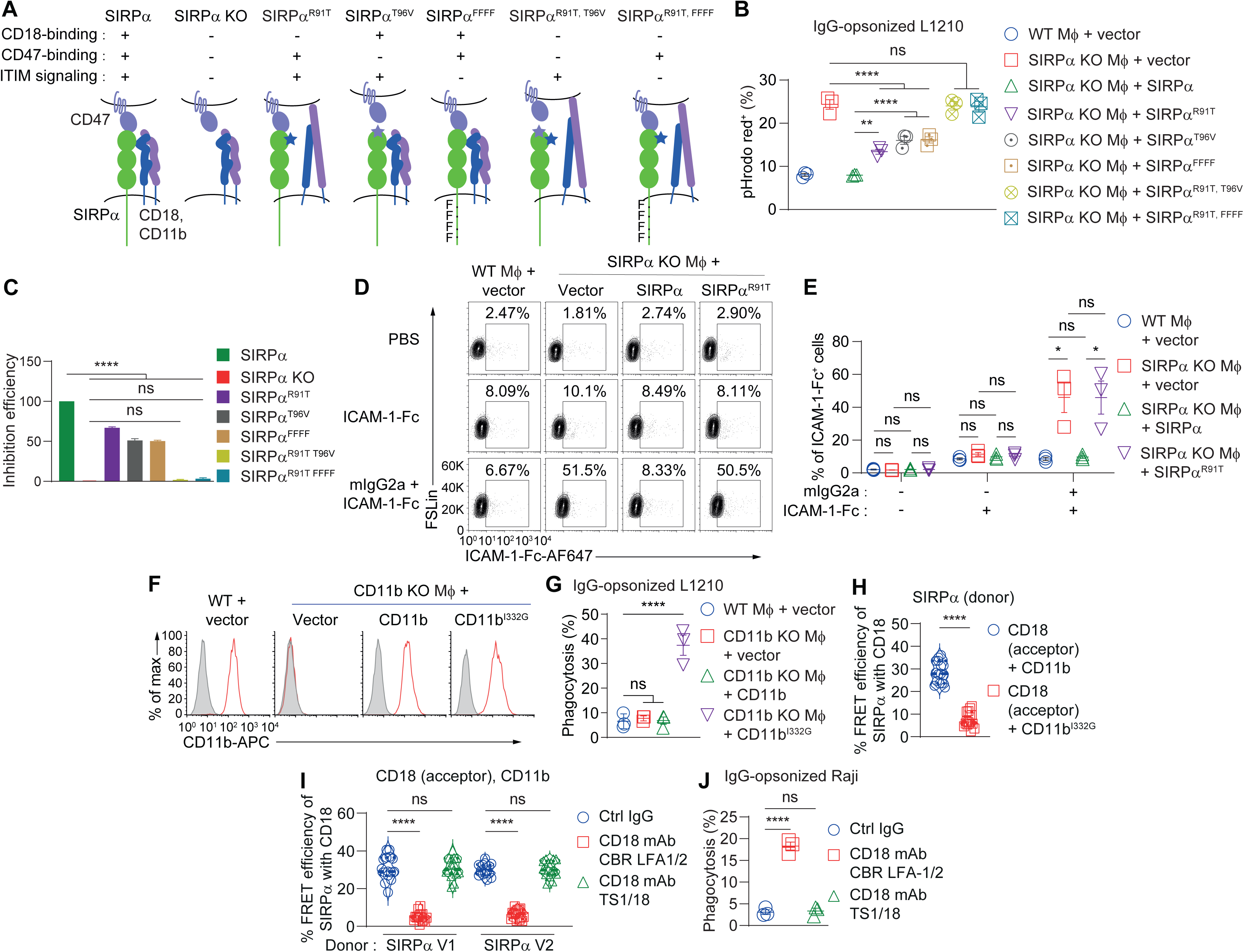
SIRPα-CD18 *cis* interaction suppresses phagocytosis and Mac-1 activation. (**A** to **C**) The impact of SIRPα variants defective in CD18-binding, CD47-binding or phosphatase signaling, alone or in combination, expressed in BMDMs, was analyzed. (**A**) Schematic depictions of SIRPα variants, as was done for Fig. 1B. SIRPα^R91T^ carried an arginine (R)-to-threonine (T) mutation at position 91 (shown by blue star), which abolished CD18-binding. (**B**) Phagocytosis assays of IgG-opsonized L1210 cells by BMDMs, as was done for Fig. 1F. (**C**) Efficiency of phagocytosis inhibition was calculated as for Fig. 1G, using values from Fig. 4B. (**D** and **E**) Representative flow cytometry profiles (**D**) and compiled data from 3 independent experiments (**E**) of ICAM-1-binding using SIRPα KO BMDMs expressing WT SIRPα or SIRPα^R91T^ BMDMs, in the presence or absence of FcR triggering using mouse IgG2a. (**F** and **G**) The impact of a SIRPα variant carrying the isoleucine-to-glycine 332 (I332G) mutation, expressed in SIRPα KO BMDMs, was analyzed. **(F)** Flow cytometry analyses of CD11b expression. (**G**) Compiled data from 3 independent phagocytosis assays, assessed by microscopy. (**H**) FRET assays of donor-labeled SIRPα, acceptor-labeled CD18 and unlabeled CD11b in the presence of WT CD11b or CD11b^I332G^, as was done for Fig. 2, D to F. (**I**) FRET assays of donor-labeled human SIRPα version (V) 1 or V2 with acceptor-labeled human CD18 and unlabeled human CD11b, in the presence of Ctrl IgG, human CD18 mAbs CBR LFA1/2 or TS1/18, as was done for Fig. 2, D to F. (**J**) Phagocytosis of human lymphoma cells Raji, which were opsonized with CD20 mAbs, by human peripheral blood monocyte (PBMC)-derived macrophages, in the presence of the indicated mAbs, was assessed by microscopy. All data are means ± s.e.m. ns, not significant; \**p* < 0.05, \*\**p* < 0.01 and \*\*\*\**p* < 0.0001. Results in (**D** and **F**) are representative of 3 independent experiments. Results in (**B**, **C**, **E** and **G** to **J**) are pooled from 3 independent experiments. Each symbol in (**B**, **E** and **G** to **J**) represents one cell, mouse or healthy donor.

In this light, we examined if the *cis* interaction prevented Mac-1 activation. Upon engagement of pro-phagocytic FcRs, but not in its absence, SIRPα KO BMDMs displayed enhanced binding to an ICAM-1 fusion protein, in keeping with Mac-1 activation, compared to WT BMDMs (**Fig. 4, D and E**). When introduced in SIRPα KO BMDMs, SIRPα^R91T^ failed to suppress binding to ICAM-1, unlike WT SIRPα. Previous studies also demonstrated that an isoleucine-to-glycine substitution at position 332 of CD11b (CD11b^I332G^), or treatment with the cation Mn²⁺, resulted in constitutive activation of Mac-1 (*15, 28*). When introduced into CD11b KO BMDMs, the CD11b^I332G^ mutant was expressed at levels comparable to WT CD11b (**Fig. 4F**). Similar to observations reported for other targets (*15, 28*), it increased phagocytosis of IgG-opsonized L1210, compared to WT CD11b (**Fig. 4G, and fig. S7A**). Interestingly, the I332G mutation also disrupted the *cis* interaction between SIRPα and CD18 (**Fig. 4h, and fig. S7B**). Mn²⁺ also increased phagocytosis—dependent on CD18—when compared to untreated cells (**fig. S7, C and D**). Moreover, the cation disrupted the *cis* interaction between SIRPα and CD18 (**fig. S7, E and F**).

Lastly, to ascertain the impact of the SIRPα-CD18 *cis* interaction in human cells, we examined the effects of human CD18 mAbs CBR LFA-1/2 and TS1/18. mAb CBR LFA-1/2 has been reported to activate Mac-1, while mAb TS1/18 did not (*29, 30*). As was seen with the mouse proteins, there was energy transfer between donor-labeled human SIRPα [either version (V)1 or V2, which both exist in humans (*31*)] and acceptor-labeled human CD18 (**Fig. 4I**). MAb CBR LFA-1/2 prevented energy transfer between human SIRPα and human CD18, whereas mAb TS1/18 did not (**Fig. 4I**). MAb CBR LFA-1/2, but not mAb TS1/18, also enhanced phagocytosis of IgG-opsonized human lymphoma cells Raji by human blood-derived macrophages (**Fig. 4J**).

Thus, the SIRPα-CD18 interaction suppressed activation of Mac-1 and phagocytosis, in synergy with the SIRPα-CD47 interaction. Activation of Mac-1 correlated with disruption of SIRPα-CD18.

## SIRPα Abs disrupting SIRPα-CD18 and SIRPα-CD47 have greater anti-tumor activity

We investigated whether SIRPα mAbs could disrupt the SIRPα-CD18 interaction, the SIRPα-CD47 interaction or both, and correlated these effects with their impact on phagocytosis. To enable these studies, as well as *in vivo* studies, we used a series of mouse SIRPα mAbs generated in our laboratory that recognized SIRPα but not SIRPβ1a, SIRPβ1b, or SIRPβ1c, apart from mAb #16, which reacted with SIRPβ1a (**fig. S8, A and B**). In FRET assays, mAb #17 demonstrated the strongest ability to diminish the SIRPα-CD18 interaction, while mAbs #23 and #27 had a small impact, compared to Ctrl IgG (**Fig. 5A**). The other mAbs, #3 and #16, showed no effect. In contrast, mAbs #23 and #27 were the most effective at blocking the binding of SIRPα to CD47 (*17*), compared to Ctrl IgG, although partial blocking effects were observed with mAbs #3, #16, and #17 (**Fig. 5B**).

**Fig. 5.**
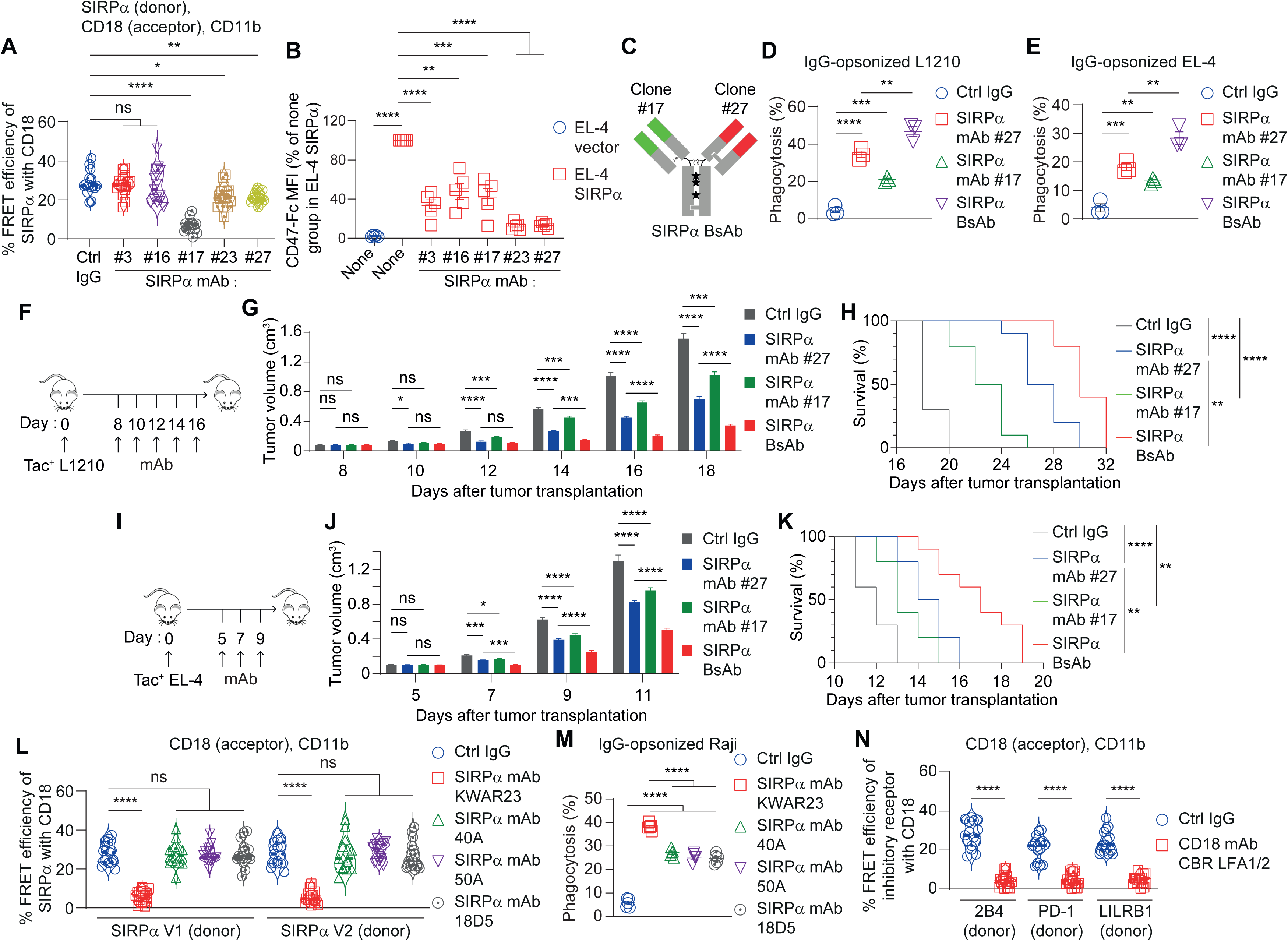
SIRPα mAbs disrupting both SIRPα-CD18 and SIRPα-CD47 have greater anti-tumor activity. (**A**) FRET assays of donor-labeled mouse SIRPα with acceptor-labeled mouse CD18 and unlabeled mouse CD11b, in the presence of Fc-silent mouse SIRPα mAbs, as was done for Fig. 2, D to F. (**B**) Binding of a soluble CD47-Fc fusion protein to EL-4 cells, expressing or not expressing mouse SIRPα, was studied by flow cytometry. (**C** to **K**) Generation and impact of bispecific antibody (BsAb) against mouse SIRPα. (**C**) Schematic representation of Fc-silent BsAb combining one arm of mAb #17 with one arm of mAb #27, using the “knob-into-hole” technology. Phagocytosis of IgG-opsonized L1210 cells (**D**) and EL-4 cells (**E**) by WT BMDMs, in the presence of mAbs, was assessed by a microscopy assays. (**F** to **K**) Schematic depictions of the assays are shown in (F and I). RAG-1 KO mice injected subcutaneously with Tac^+^ L1210 cells (**G** and **H**), or C57BL/6J mice injected subcutaneously with Tac^+^ EL-4 cells (**J** and **K**), were treated by intraperitoneal injection of Fc-silent mAbs, alongside Tac mAb 7G7 for opsonization. Tumor volume was measured using a caliper (**G** and **J**) and survival was recorded (**H** and **K**). (**L**) FRET assays of donor-labeled human SIRPα V1 or V2 with acceptor-labeled human CD18 and unlabeled human CD11b in the presence of Fc-silent Ctrl IgG and human SIRPα mAbs KWAR23, 40A, 50A, or 18D5, as was done for Fig. 2, D to F. The mAbs were rendered Fc-silent by the LALAPG mutation. (**M**) Phagocytosis of IgG-opsonized Raji cells by human macrophages in the presence of Fc-silent Ctrl IgG and SIRPα mAbs KWAR23, 40A, 50A, or 18D5, was assayed as for Fig 4J. (**N**) FRET assays of donor-labeled human 2B4 (SLAMF4), PD-1 or LILRB1 with acceptor-labeled human CD18, in the presence of Ctrl IgG or human CD18 mAb were done as for Fig. 2, D to F. All data are means ± s.e.m. ns, not significant; \**p* < 0.05, \*\**p* < 0.01, \*\*\**p* < 0.001 and \*\*\*\**p* < 0.0001. Results are pooled from a total of two (**H** and **K**), three (**A**, **D**, **E**, **G**, **J**, **L** and **N**) or five (**B** and **M**) independent experiments. Each symbol in (**A**, **D**, **E** and **L** to **N**) represents one healthy donor, cell or mouse.

For maximal blockade of SIRPα, we generated a bispecific antibody (BsAb) that combined one arm of mAb #17 with one arm of mAb #27, using the “knob-into-hole” technology (*32*) (**Fig. 5C**). This BsAb robustly inhibited the SIRPα-CD18 *cis* interaction and the SIRPα-CD47 *trans* interaction, compared to Ctrl IgG (**fig. S8, C to E**). It was also more effective than mAb #17 or mAb #27 at enhancing the phagocytosis of IgG-opsonized L1210 or EL-4 cells by WT BMDMs, compared to Ctrl IgG (**Fig. 5, D and E**). Agreeing with these data, in BMDMs expressing SIRPα^R91T^, the BsAb was as effective as mAb #27 at promoting phagocytosis, whereas in BMDMs containing SIRPα^T96V^, the BsAb was like mAb #17 (**fig. S8F**).

To evaluate the anti-tumor efficacy of the BsAb, we first utilized RAG-1-deficient mice, which lack T cells and B cells, that were inoculated subcutaneously with Tac-positive L1210 cells. Once tumor growth was established, mice were treated with the BsAb or the mAbs, alongside the Fc-active Tac mAb 7G7 for opsonization. Compared to mAb #17 or mAb #27, the BsAb resulted in greater inhibition of tumor growth and improved mouse survival, when contrasted to Ctrl IgG (**Fig. 5, F to H, and fig. S9A**). Similar results were noted when EL-4 cells were inoculated into immunocompetent syngeneic C57BL/6J mice (**Fig. 5, I to K, and fig. S9B**).

All therapeutic human SIRPα mAbs reported to date have been developed with the aim of disrupting the SIRPα-CD47 interaction. We uncovered that MAb KWAR23, but not mAbs 40A, 50A, or 18D5, also blocked the SIRPα-CD18 *cis* interaction in FRET assays, compared to Ctrl IgG (**Fig. 5L**). This blocking effect was validated in PLA assays (**fig. S10, A and B**). All SIRPα mAbs inhibited binding of CD47-Fc to SIRPα compared to Ctrl IgG, although this effect was limited to the V1 isoform for mAb 18D5, as previously reported (*33*) (**fig. S10C**). Consistent with its dual blocking activity towards SIRPα, mAb KWAR23 was more effective than the other SIRPα mAbs at enhancing phagocytosis of IgG-opsonized Raji cells by human macrophages, compared to Ctrl IgG (**Fig. 5M**). One caveat with these SIRPα mAbs is that all also reacted with other SIRP family members, including SIRPβ1, SIRPβ2 and SIRPγ (**fig. 10D**).

Therefore, SIRPα Abs disrupting the SIRPα-CD47 and SIRPα-CD18 interactions, such as a SIRPα BsAb and the “bifunctional” human mAb KWAR23, caused a greater increase in phagocytosis and better tumor suppression, compared to mAbs that primarily interrupted either interaction (**fig. S11**).

## Other inhibitory checkpoint receptors interact with CD18

As CD18 plays a critical role in activation of multiple types of immune cells, we speculated that other inhibitory receptors may interact in *cis* with CD18. To test this notion, FRET assays were performed using acceptor-labeled human CD18 and donor-labeled human PD-1, which inhibits T cell activation, and 2B4 or LILRB1, which inhibit phagocytosis. All donors displayed energy transfer to CD18, and this effect was prevented by blocking CD18 mAb CBR LFA1/2 (**Fig. 5N**). Thus, multiple inhibitory receptors interacted in *cis* with CD18, suggesting that this mechanism of inhibition was shared by other checkpoints.

## Discussion

Considerable efforts have been directed toward identifying additional inhibitory immune checkpoints, such as SIRPα-CD47 in macrophages, as targets for cancer immunotherapy, although these strategies have typically led to mitigated results in the clinic (*34*). In our study, we found that blockade of the SIRPα-CD47 *trans* interaction, which has been targeted by the existing SIRPα-CD47 blockade therapies, only partially eliminated the inhibitory function of SIRPα, possibly explaining the limited efficacy of these agents in the clinic. Another study had also suggested a CD47-independent role of SIRPα, although the mechanism was not clarified (*35*).

This phenomenon was attributed to an additional mechanism of suppression by SIRPα. SIRPα interacted in *cis* with CD18 at the macrophage plasma membrane, thereby hindering Mac-1 activation, phagocytosis and anti-tumor immunity. This *cis* interaction required SIRPα residues, particularly R91, that were distinct from those involved in the SIRPα-CD47 *trans* interaction. Although the nearby residue T96 of SIRPα is directly implicated in the interaction with CD47 molecular (*16*), we do not know if R91 was directly binding to CD18 or was generating a SIRPα conformation needed for CD18-binding. Although others reported some biochemical evidence of an interaction between SIRPα and Mac-1, this interaction involved CD11b rather than CD18 and occurred in *trans* (*36*).

From a physiological perspective, the *cis* mechanism of inhibition may be crucial for preventing Mac-1 activation in response to weak stimuli, which could lead to immunopathology if unopposed, especially considering that Mac-1 has many ligands including some that are broadly expressed on normal cells and soluble ones (*37*). Interestingly, however, strong activation of Mac-1 by mutation of CD11b or Mn^2+^ led to dissociation of CD18 from SIRPα, suggesting that once a threshold is reached, activation of Mac-1 is sustained by loss of the *cis* interaction. From a therapeutic standpoint, these findings suggested that dual blockade of the SIRPα-CD18 *cis* interaction and the SIRPα-CD47 *trans* interaction—in a manner achieved by our newly engineered SIRPα BsAb or SIRPα mAb KWAR23—may be necessary to maximize phagocytosis and anti-tumor immunity in clinical settings. Although clinical trials involving mAb KWAR23 have not yet been reported, this antibody also blocked the interaction between CD47 and SIRPγ, an activating receptor on T cells, which may limit its clinical value (*38, 39*). Finally, it should be pointed out that a similar *cis* interaction targeting CD18 was noted for other inhibitory receptors, such as PD-1, suggesting that this supplementary inhibitory mechanism applies to other inhibitory checkpoints and should be broadly considered in drug design.

## Methods

### Mice

Mice lacking SIRPα (*Sirpa^-/-^*) in the C57BL/6J background were generated in our laboratory, as described (*17*). CD18 knock-out (KO) (*Itgb2*^-/-^), RAG-1 KO (*Rag1^-/-^*) and C57BL/6J mice were obtained from Jackson Laboratory (Bar Harbor, ME). CD45 KO (*Ptprc^L3X/L3X^*) mice, which lack expression of CD45, were obtained from Dr. Silvia Vidal (McGill University, Montreal, Canada) (*40*). SIRPα KO mice were bred with CD45 KO or CD18 KO mice to create SIRPα-CD45 double (D) KO mice and SIRPα-CD18 DKO mice. All mice were maintained in the C57BL/6J background. They were kept in a specific pathogen-free environment. Either males or females were used, between 8 to 12 weeks of age. Littermates were used as control in most experiments, except for some studies where wild-type syngeneic, age and sex-matched mice were used. Animal experimentation was done in accordance with the Canadian Council of Animal Care and approved by the IRCM Animal Care Committee.

### Cells

Macrophages were generated as previously detailed (*13, 17*). In summary, mouse bone-marrow-derived macrophages (BMDMs) were obtained by culturing freshly isolated bone marrow cells in a medium enriched with 30% (vol/vol) L929 cell-conditioned medium, providing colony-stimulating factor-1 (CSF-1) for a period of 7 d. Human macrophages were derived from peripheral blood mononuclear cells isolated of healthy donors via Ficoll-Paque PLUS (Cat# 36-101-6383, GE Healthcare, Marlborough, MA), with approval from the IRCM Human Ethics Board. The isolated peripheral blood mononuclear cells were seeded into Petri dishes containing serum-free RPMI medium and allowed to adhere for 30 min. Following gentle washes to eliminate non-adherent cells, the adherent cells—which predominantly consist of monocytes—were differentiated into macrophages by incubating them in a medium supplemented with 10% human serum and 10 ng ml^-1^ CSF-1 (Cat# 300-25, Peprotech, Rocky Hill, NJ) for 7 d. The cell lines L1210 (CCL-219), P815 (TIB-64), EL-4 (TIB-39), L929 (CCL-1), HEK293T (CRL-3216), Phoenix-Eco (CRL-3214), and Raji (CCL-86) were sourced from the American Type Culture Collection (Rockville, MD). The provider authenticated the cells, and the identity of immune cell lines was confirmed by flow cytometry. All cells tested negative for Mycoplasma.

### cDNAs, mutants and retroviral infection

Mouse cDNAs for SIRPα, SLAMF7, CD18, CD11a and CD11b were described(*13, 14, 22*). Mouse cDNAs encoding SIRPβ1a (Cat: OMu10495) or SIRPβ1c (Cat: OMu31303) was from GenScript (Pistacaway, NJ). Mouse SIRPβ1b cDNA (Cat: SKU MG219500) was from OriGene (Rockville, MD). Human cDNAs for SIRPα version (V) 1 (Cat: HG11612-UT), PD-1 (Cat: HG10377-M), LILRB1 (Cat: HG16014-UT), 2B4 (Cat: HG10042-NF), CD18 (Cat: HG10970-UT) and CD11b (Cat: HG10494-UT) were obtained from Sino Biological (Beijing, China). Human SIRPα V2 cDNA (Cat: SC125649) was from OriGene (Rockville, MD). Human cDNAs encoding SIRPβ1-Fc (Cat: #156780), SIRPβ2-Fc (Cat: #156705) and SIRPγ-Fc (Cat: #156772) were obtained from Addgene (Cambridge, MA).

The following mutated cDNAs were generated by overlap extension PCR. Mouse SIRPα: SIRPα IgV domain, which lacked the second and third Ig-like domain; SIRPα^FFFF^, which had tyrosine (Y)-to-phenylalanine (F) substitutions at Y436, 460, 477, and 501; SIRPα^ΔIC^, which was truncated after arginine 401 in the cytoplasmic domain; SIRPα^T96V^; SIRPα^R91T^; SIRPα^T96V, FFFF^; _SIRPa_T96V, ΔIC; _SIRPα_R91T, T96V; _SIRPα_R91T, FFFF; _SIRPα_S81P, A83T, Y86H, V87S, R91T; _SIRPα_T37I, E40V, V43F, S44F, A46G; _SIRPα_A116S, I118T; _SIRPα_T133I, V143L, Y144L; _SIRPα_S81P_, SIRPα_A83T_, SIRPα_Y86H_, SIRPα_V87S and SIRPα^R91T^. Mouse CD11b: CD11b^I332G^ carrying a isoleucine (I)-to-glycine (G) mutation at position 332.

Wild-type or mutated cDNAs were cloned into the pFB-GFP or the pMIGR-GFP retroviral vector, which also encode the green fluorescent protein (GFP). After transfection in Phoenix-Eco cells, viral supernatants were recovered and used for spin infection of WT BMDMs, SIRPα KO BMDMs, P815 or EL-4 cells. Plasmids expressing GFP alone were used as controls. 48 h after retroviral infection, GFP-positive cells were sorted and used for experimentation. P815 and EL-4 cells derivatives expressing the Tac antigen (human CD25) or mouse SIRPα, were sorted a second time using fluorescence-labeled mAbs. Tac-positive L1210 derivatives were described (*13, 17*).

### Antibodies

Mouse SIRPα mAbs #3, #16, #17, #23 and #27 were generated as reported (*17*). Mouse SIRPα mAb MY-1 was provided by Takashi Matozaki (Kobe University, Kobe, Japan). Mouse CD18 mAb M18/2 was purchased from Developmental Studies Hybridoma Bank (Iowa City, IA). To create recombinant versions of mAbs, cDNA sequences encoding the variable regions of heavy chain (VH) and light chain (VL) were determined using RNA sequencing, as outlined (*17*). In all cases, VH-and VL-encoding cDNAs were cloned in-frame into one or both of the following expression plasmids (*17*): pAb-mIgG2a(LALAPG), which contains genes encoding Fc-silent mouse IgG2a (mIgG2a) with the LALAPG mutation, and pAb-mIgK; or pAb-hIgG1(LALAPG), which contains genes encoding Fc-silent human IgG1 with the LALAPG mutation, and pAb-hIgK. Recombinant mAbs were expressed in HEK293T or CHO cells and purified with protein A Sepharose (Cat# GE17-1279-03, Sigma-Aldrich, St. Louis, MO).

Mouse SIRPα bispecific antibody (BsAb) was generated using the “knob-into-hole” technology (*32, 41*). The VH domain of SIRPα mAb #17 was cloned into pAb-hIgG1(LALAPG; hole), which encoded in-frame the Fc portion of human IgG1 with a “LALAPG” mutation (leucine 234-to-alanine 234; leucine 235-to-alanine 235; proline 329-to-glycine 329 mutation that prevents binding to FcRs), a “T366S, L368A, Y407V” mutation (threonine 366-to-serine 366, leucine 368-to-alanine 368, tyrosine 407-to-valine 407) that creates a “hole” in the Fc portion, and a “H435R, Y436F” mutation (histidine 435-to-arginine 435, tyrosine 436-to-phenylalanine 436) that prevents hole-hole homodimers; the VL domain of mAb #17 was cloned into pAb-hIgK. Conversely, the VH domain of SIRPα mAb #27 was cloned into pAb-hIgG1(LALAPG; knob), which encodes in-frame the Fc portion of human IgG1 carrying a “LALAPG” mutation, a “T366W” mutation (threonine 366-to-tryptophan 366) that makes a “knob” in the Fc portion, a “F126C” mutation (phenylalanine 126-to-cysteine 126) that creates a new cysteine for binding to the mutant light chain, and a “C220V” mutation (cysteine 220-to-valine 220) that removes the original cysteine; the VL domain of mAb #27 was cloned into pAb-hIgK with a “S121C” mutation (serine 121-to-cysteine 121) that makes a new cysteine for binding to the mutant heavy chain and a “C214V” mutation (cysteine 214-to-valine 214) that removes the original cysteine. To generate Fab fragments of human SIRPα mAb KWAR 23 and control IgG MOPC-21, the VH domain of these mAbs was cloned into pAb-hIgG1(Fab), which encodes in-frame the Fc portion of a truncated human IgG1 (ending with lysine 222) and is followed by a glycine-serine (GS) linker and an 8x His tag. The VL domain was cloned into pAb-hIgK.

All recombinant mAbs were quantified by SDS-PAGE and Coomassie blue staining, using bovine serum albumin (BSA) as a standard.

For opsonization of target cells with IgG, Fc-intact human CD25 mAb 7G7 (mouse IgG2a, BioXCell, Lebanon, NH) and Fc-intact human CD20 mAb rituximab biosimilar (human IgG1, BioXCell) were used. For flow cytometry assays, CD45 mAb 30-F11, CD11b mAb M1/70, F4/80 mAb BM8, Ly6G mAb 1A8, NK1.1 mAb PK136, TCR mAb H57-597, CD8 mAb 53-6.7, CD4 mAb RM4-5, SIRPα mAb P84 and CD18 mAb M18/2 were obtained from Biolegend (San Diego, CA). For immunoprecipitations and immunoblots, we utilized SIRPα Ab (Cat# PA5-19869, Thermo Fisher Scientific, Waltham, MA), CD18 mAb C71/16 (Santa Cruz Biotechnologies, Dallas, TX), CD11b mAb EPR1344 (Abcam, Ontario, Canada) and SIRPα rabbit antiserum (generated in our laboratory) (*14*).

### Fusion proteins

To produce Fc fusion proteins containing the extracellular domain of mouse CD47, mouse ICAM-1, mouse SIRPα, mouse SIRPβ1a, mouse SIRPβ1b, mouse SIRPβ1c, human SIRPα V1 or human SIRPα V2, cDNAs were cloned into pFc-hIgG1(LALAPG). For production, cDNAs were transfected in HEK293T or CHO cells. Then, fusion proteins were purified with protein A Sepharose (Cat# GE17-1279-03, Sigma-Aldrich, St. Louis, MO). A histidine (His)-tagged version of mouse CD47 (CD47-His; Cat# 57231-M08H) was from Sino Biological (Beijing, China).

To produce His-tagged mouse SIRPα, mouse SIRPα^R91T^, mouse CD11b and mouse CD18, we generated cDNAs encoding the extracellular segment of these proteins, in which a SNAP tag was inserted after the amino-terminal signal peptide and a 10x His tag was inserted at the carboxyl-terminal end of the extracellular domain. cDNAs were cloned in the pPPI4 vector. After transfection in HEK293T or CHO cells, recombinant proteins were purified from cell culture medium using GE Ni Sepharose® Excel (GE17371201, Sigma-Aldrich, St. Louis, MO) and eluted with 0.5 M imidazole. Subsequently, His-tagged proteins were labeled with SNAP-Cell-505 (Cat# S9103S, New England Biolabs, Ipswich, MA) or SNAP-Cell-TMR (Cat# S9105S, New England Biolabs, Ipswich, MA) following the manufacturer’s instructions. Free dye was removed using a PD-10 desalting column (Cat# 87766, Thermo Fisher Scientific, Waltham, MA). All proteins were quantified by SDS-PAGE and Coomassie blue staining, using BSA as a standard.

### Flow cytometry and Fc fusion protein-binding assay

Cells were harvested and washed with 2% (vol/vol) fetal bovine serum (FBS)-containing phosphate-buffered saline (PBS). To prevent non-specific binding to FcRs, FcRs were blocked with a mix of mouse IgG2a (mAb 7G7) and mouse CD16/32 mAb 2.4G2 for 30 min on ice. Afterwards, cells were stained with the indicated fluorophore-conjugated antibodies for 30 min on ice. After being washed, cells were analyzed using flow cytometer. To test binding of CD47-Fc and ICAM-1-Fc to BMDMs, cells were incubated for 30 min on ice with Fc-silent fusion proteins, washed and incubated for 30 min on ice with Alexa Fluor 647-labeled F(ab′)_2_ fragments goat anti-human IgG1, Fc-specific (Cat# 109-606-098, Jackson Immune Research, Bar Harbor, ME). After additional washes, fluorescence was evaluated by flow cytometry. To examine the impact of SIRPα Abs on CD47-Fc binding, EL-4 cells expressing or not expressing mouse or human SIRPα variants were incubated for 30 min on ice with the indicated Fc-silent Abs.

Subsequently, cells were stained with Alexa Fluor 647-labeled Fc-silent CD47-Fc (generated in our laboratory) for 30 min on ice. After washes, fluorescence was evaluated by flow cytometry. To study the competition between CD18 and CD47 for binding to SIRPα, HEK293T cells were transfected with the indicated plasmids for 48 h before staining with fluorescence-labeled fusion proteins or mAbs for 30 min on ice. After washes, fluorescence was evaluated by flow cytometry.

### Immunoprecipitations, immunoblots and mass spectrometry

BMDMs from WT or SIRPα KO mice were lysed in Brij99-containing buffer, and were subjected to immunoprecipitation, immunoblot or mass spectrometry, as reported (*17, 42*). The following criteria were used to select potentially relevant SIRPα interactors: 1) presence in SIRPα immunoprecipitates from WT BMDMs, but not from SIRPα KO BMDMs; 2) observation in three independent experiments.

### Phagocytosis assays

Phagocytosis was evaluated using a pHrodo-based assay or a microscopy-based assay, as reported (*13, 17*). Tac-positive mouse target cells L1210, P815 or EL-4 were opsonized with Tac mAb 7G7 (a mouse IgG2a), whereas human target cell Raji was opsonized with human CD20 mAb rituximab biosimilar (a human IgG1).

### Sub-cutaneous tumor transplantation assay

Tac-positive L1210 and EL-4 derivatives (0.5×10^6^ cells) were injected subcutaneously into the right flank of 8-to 10-week-old RAG-1 KO or wide-type C57BL/6J, respectively. When tumor size was reached 5×5 mm^2^ in diameter, mice were injected intraperitoneally with 200 μg of Fc-silent control IgG mAb MOPC-21, SIRPα mAb #27, SIRPα mAb #17 or SIRPα BsAb, alongside 200 μg of Fc-intact Tac mAb 7G7 for opsonization. Experiments were terminated when or before tumor volume reached 1.5 cm^3^. Tumors were then dissected, weighed and processed as described.

### Fluorescence resonance energy transfer assay

For the fluorescence resonance energy transfer (FRET) assay, constructs encoding CLIP-tagged SIRPα variants, SIRPβ1a, SLAMF7, 2B4, PD-1 or LILRB1 were co-transfected with constructs encoding SNAP-tagged CD18 or CD11b into HEK293T cells. Untagged version of CD18, CD11b or CD11a were also added for stable plasma membrane expression of the integrin complexes. After 2 d, cells were collected and seeded onto poly-D-lysine (Cat# P6407, Sigma-Aldrich, St. Louis, MO)-treated confocal microscopy plates. The next day, cells were labeled with CLIP-Surface 547 (Cat# S9233S, New England Biolabs, Ipswich, MA) and SNAP-Surface Alexa Fluor 647 (Cat# S9136S, New England Biolabs, Ipswich, MA) for 45 min at 37°C, in the presence or absence of 10 μg ml^-1^ of mAbs. Cells were then fixed with 4% paraformaldehyde (Cat# 22023, Biotium, CA) and washed, prior to the FRET assay. Images were acquired with an LSM710 confocal microscope (Carl Zeiss) by exciting CLIP-Surface 547 (energy donor) at 543 nm and SNAP-Surface Alexa Fluor 647 (energy acceptor) at 635 nm. Images before and after acceptor bleaching were acquired for FRET analysis, using ImageJ (Fiji) with the AccPbFRET plugin, as previously described (*17*).

### Large unilamellar vesicle reconstitution assay

Large unilamellar vesicle (LUV) reconstitution followed by FRET assay was performed as described (*17*). For LUV preparation, 80% 1-palmitoyl-2-oleoyl-glycero-3-phosphocholine (POPC, Cat# 850457C, Avanti Polar Lipids, Alabaster, AL) and 20% 1,2-dioleoyl-sn-glycero-3-[(N-(5-amino-1-carboxypentyl)iminodiacetic acid)succinyl] (nickel salt) (DGS-NTA-Ni, Cat# 790404C, Avanti Polar Lipids, Alabaster, AL) were mixed in chloroform, dried under a stream of nitrogen, desiccated for 1 h in a vacuum container and resuspended in PBS. LUVs were generated by extrusion 20 times through a pair of polycarbonate filters containing pores of 200 nm diameter.

8.3 nM SNAP-Cell-505-labeled mouse SIRPα-His or mouse SIRPα^R91T^-His was mixed with 0.23 nM LUVs harboring DGS-NTA-Ni in PBS containing 1.5 mg ml^-1^ BSA and 1 mM tris(2-carboxyethyl)phosphine (TCEP, Cat: T2556, Thermo Fisher Scientific, Waltham, MA) in a 96-well solid black microplate, during which the SNAP-Cell-505 fluorescence was monitored in real time, using a plate reader with 504-nm excitation and 540-nm emission. After 20 min, the fluorescence reading was paused, and the SNAP-Cell-TMR-labeled second protein component, that is 25 nM mouse CD18-His or mouse CD11b-His, was injected and the fluorescence was monitored for 20 min. Data were normalized for the mean fluorescence intensity of the last 10 data points, before the addition of SNAP-Cell-TMR-labeled protein and plotted with GraphPad Prism.

### Proximity ligation assay

Proximity ligation assay (PLA) was performed with a commercial kit according to the manufacturer’s recommendation (Cat# DUO92008, Duolink In Situ, Sigma-Aldrich, St. Louis, MO) (*17*). Briefly, mouse or human macrophages were incubated in the presence or in the absence of CD47-His or Fab fragments of mAb KWAR23 (10 μg ml^-1^) for 30 min, respectively, and then fixed to stabilize proximity. Fab fragments of mAb KWAR23 were used to prevent recognition by the secondary mAb used in the PLA. Fc-silent CD18 mAb M18/2 (a human IgG1) and SIRPα mAb #27 (a mouse IgG2a) were then added to mouse BMDMs, while Fc-silent CD18 mAb TS1/18 (a mouse IgG1) and SIRPα mAb 50A (a human IgG1) were added to human macrophages, followed by oligonucleotide-linked secondary antibodies specifically recognizing the Fc portion of each of the primary antibodies. Then, proximity was detected by PCR with complementary oligonucleotides coupled to fluorochromes, and a LSM710 confocal microscope (Carl Zeiss). In each experiment, five images per condition were randomly chosen. A total of 60 cells from three independent experiments were randomly chosen for quantification

## Statistical analyses

Prism 9 software (GraphPad) was used for paired or unpaired Student’s *t* tests (two-tailed), and for one-way ANOVA followed by Tukey’s or Dunnett’s multiple comparison tests, when appropriate. The normal distribution of the data was tested using the D’Agostino-Pearson normality test (Prism 9) and parametric tests used for statistical analyses accordingly.

## Data availability

All relevant data are available from the corresponding author upon reasonable request.

## Acknowledgements

We thank the members of the Veillette laboratory for useful discussions. This work was supported by grants from the Canadian Institutes of Health Research (MT-14429, MOP-82906, FDN-143338, PJT-178314 and PJT-183593), the Terry Fox Research Institute (1190–02) and the Ministère de l’économie et de l’innovation (MEI; Québec) to A.V.; SRG2024-00027-FHS and UMDF-TISF/2025/002/FHS to Z.T. Z.T. received a Fellowship from the Cole Foundation; and A.V. held the Canada Research Chair on Signaling in the Immune System.

## Contributions

Z.T., M.C.-Z., J.Q., L.S.W., J.L., D.D. and A.V. designed experiments. Z.T., M.C.-Z. and J.Q. performed experiments. All authors interpreted the results. Z.T. and A.V. wrote the manuscript. All authors commented on the manuscript.

## Figure legends

**fig. S1. SIRPα inhibits phagocytosis independently of CD47-binding and phosphatase signaling.** (**A**) SIRPα has 1 IgV and 2 IgC domains in its extracellular component, a single transmembrane domain, and a cytoplasmic domain with 4 ITIMs. The amino acid sequence of the transmembrane and cytoplasmic domains of SIRPα from the C57BL/6 mouse, as well as the location of key mutations, are displayed. Amino acid numbering in based on full-length including the leader peptide. V, variable. C, constant. (**B** and **C**) Same as Fig. 1, E and F, except that no target cells were added in the assays. (**D**) Compiled data from phagocytosis assays of IgG-opsonized L1210, P815 and EL-4 cells by BMDMs expressing SIRPα variants as in Fig. 4G. All data are means ± s.e.m. ns, not significant; \*\*\**p* < 0.001 and \*\*\*\**p* < 0.0001. Results in (**B**) are representative of 6 independent experiments, except for SIRPα^T96V^, SIRPα^T96V, FFFF^ and SIRPα^T96V, ΔIC^ that are representative of 3 experiments. Results in (**C**) are pooled from 6 experiments, except for SIRPα^T96V^, SIRPα^T96V, FFFF^ and SIRPα^T96V, ΔIC^ that were from 3 experiments, whereas results from (**D**) are from 3 independent experiments, except for vector, WT SIRPα and SIRPα^FFFF^ with IgG-opsonized L1210 cells that are from 6 different experiments. Each symbol in (**C** and **D**) represents one mouse.

**fig. S2. Mac-1, not CD45, is necessary for enhanced phagocytosis in the absence of SIRPα.** (**A**) Means of normalized total ion current (TIC) for the potential interactors with SIRPα. Full names and functions are also listed. (**B** and **C**) Representative flow cytometry profiles (**B**) and compiled data from multiple phagocytosis assays (**C**) of IgG-opsonized L1210 cells by BMDMs. DKO, double KO. (**D** and **E**) Compiled data from multiple phagocytosis assays of IgG-opsonized L1210 cells by BMDMs, in the presence of CD18 mAb GAME-46 (**E**) or CD11b mAb 5C6 (**D**). (**F**) Compiled data from multiple phagocytosis assays of IgG-opsonized L1210 cells by BMDMs. All data are means ± s.e.m. ns, not significant; \*\**p* < 0.01, \*\*\**p* < 0.001 and \*\*\*\**p* < 0.0001. Results in (**B**) are representative of 3 independent experiments. Results are pooled from a total of 3 (**C** to **E**) or 4 (**F**) independent experiments. Each symbol in (**C** to **F**) represents one mouse.

**fig. S3. SIRPα interacts with CD18, while SLAMF7 interacts with CD11b.** (**A** and **B**) Representative confocal microscopy images (**A**) and compiled data from 3 independent FRET assays (**B**) using donor-labeled SIRPα, acceptor-labeled CD11b and unlabeled CD18, as was done for Fig. 2, D to F. (**C** and **D**) Representative confocal microscopy images (**C**) and compiled data from 3 independent FRET assays (**D**) using donor-labeled SIRPα, acceptor-labeled CD18 and unlabeled CD11a, as was done for Fig. 2, D to F. (**E** and **F**) Representative confocal microscopy images (**E**) and compiled data from three independent FRET assays (**F**) of donor-labeled SLAMF7, acceptor-labeled CD11b and unlabeled CD18, as was done for Fig. 2, D to F. (**G**) Model of the interaction between SIRPα, CD18, CD11b and SLAMF7. All data are means ± s.e.m. ns, not significant; \*\*\*\**p* < 0.0001. Results in (**A**, **C** and **E**) are representative of 3 independent experiments. Results in (**B**, **D** and **F**) are pooled from a total of 3 independent experiments. Each symbol in (**B**, **D** and **F**) represents one cell.

**fig. S4. Identification of amino acids critical for SIRPα-CD18 *cis* interaction.** (**A**) Amino acid sequence of the IgV domain of mouse SIRPα and SIRPβ1a, including the signal peptide. Identical residues are indicated by asterisks, whereas conserved and semi-conserved residues are shown by colons and periods, respectively. (**B** to **F**) Representative confocal microscopy images of Fig. 3B (**B**), Fig. 3D (**C**), Fig. 3F (**D**) and Fig. 3H (**E** and **F**). (**G**) LUV-based FRET assay of donor-labeled SIRPα or SIRPα^R60T^ with acceptor-labeled CD18 was monitored, as was done for Fig. 2, G and H. Results in (**B** to **G**) are representative of 3 independent experiments.

**fig. S5. SIRPα utilizes different epitopes for binding to CD18 and CD47.** (**A** to **C**) Representative flow cytometry profiles of CD47-Fc binding (**A**) and pooled data from multiple experiments (**B**) with HEK293T cells expressing or not WT SIRPα or SIRPα^R60T^. Staining with SIRPα mAb P84 staining is shown in (**C**). (**D** and **E**) Representative confocal microscopy images (**D**) and compiled data from 3 independent experiments (**E**) of PLA for SIRPα and CD18, in the presence or absence of CD47-His fusion protein. Scale bar, 10 μm. (**F** to **I**) Representative flow cytometry profiles of 3 independent experiments of CD47-Fc-binding (**F**), and CD18 mAb (**H**) or SIRPα mAb staining (**I**) with HEK293T cells expressing or not mouse SIRPα and Mac-1; (**G**) was compiled data from 3 independent experiments of (**F**). (**J** to **L**) Representative flow cytometry profiles of 3 independent experiments of CD47-Fc-binding (**J**) and CD18 staining (**L**) with WT or CD18 KO BMDMs; (**K**) was compiled data from 3 independent experiments of (**J**). All data are means ± s.e.m. ns, not significant; \**p* < 0.05 and \*\**p* < 0.01. Results in (**A**, **C**, **D**, **F**, **H**, **I**, **J** and **L**) are representative of 3 independent experiments. Results in (**B**, **E**, **G** and **K**) are pooled from 3 independent experiments. Each symbol in (**E** and **K**) represents one mouse.

**fig. S6. Disruption of SIRPα-CD18 *cis* interaction increases phagocytosis.** (**A** to **D**) Same as Fig. 4, A to B. (**A** to **C**) Representative flow cytometry profiles. (D) Compiled data of 3 independent experiments from (**C**). (**E**) Compiled data from different phagocytosis assays of IgG-opsonized L1210, P815 and EL-4 cells by BMDMs expressing different SIRPα variants, as done for Fig. 4G. All data are means ± s.e.m. ns, not significant; \*\*\*\**p* < 0.0001. Results in (**A** to **C**) are representative of 3 independent experiments. Results in (**D** and **E**) are pooled from 3 independent experiments, except for (**E**, IgG-opsonized L1210 cells) that are pooled from 5 independent experiments. Each symbol in (**D** and **E**) represents one mouse.

**fig. S7. Activation of Mac-1 is paralleled by dissociation of SIRPα-CD18 *cis* interaction to promote phagocytosis.** (**A**) Same as Fig. 4G, except that parental L1210 cells were used as targets. (**B**) Compiled data from 3 independent FRET assays with donor-labeled SIRPα, acceptor-labeled WT CD11b or CD11b^I332G^ and unlabeled CD18, as was done for Fig. 2, D to F. (**C** and **D**) Compiled data from 3 independent phagocytosis assays of IgG-opsonized L1210 cells (**C**) or non-opsonized L1210 cells (**D**) by WT BMDMs. (**E** and **F**) Compiled data from 3 independent FRET assays with donor-labeled SIRPα, acceptor-labeled CD18 and unlabeled CD11b (**E**) or donor-labeled SIRPα, acceptor-labeled CD11b and unlabeled CD18 (**F**), as was done for Fig. 2, D to F. All data are means ± s.e.m. ns, not significant; \*\**p* < 0.01, \*\*\**p* < 0.001 and \*\*\*\**p* < 0.0001. Results in (**A** to **F**) are pooled from 3 independent experiments. Each symbol in (**A** to **F**) represents one mouse or cell.

**fig. S8. Characterization of mouse SIRPα antibodies.** (**A**) Representative flow cytometry profiles of EL-4 cells expressing or not full-length mouse SIRPα or IgV-only mouse SIRPα. (**B**) Compiled data from 3 independent binding assays with mouse SIRPα mAbs and the indicated fusion proteins, using ELISA. (**C**) Compiled data from 3 independent FRET assays of mouse SIRPα with mouse CD18, in the presence of Ctrl IgG or mouse SIRPα BsAb, as was done for Fig. 2, D to F. (**D** and **E**) Representative flow cytometry profiles (**D**) and compiled data from 3 independent experiments (**E**) of CD47-Fc binding to EL-4 cells expressing SIRPα, in the presence of Ctrl IgG or SIRPα BsAb. (**F**) Compiled data from different phagocytosis assays of IgG-opsonized L1210 cells by BMDMs expressing different SIRPα variants in the presence of mouse SIRPα antibodies, as done for Fig. 4G. All data are means ± s.e.m. ns, not significant; \**p* < 0.05, \*\**p* < 0.01, \*\*\**p* < 0.001 and \*\*\*\**p* < 0.0001. Results in (**A** and **D**) are representative of 3 independent experiments. Results in (**B**, **C**, **E** and **F**) are pooled from 3 independent experiments. Each symbol in (**C** and **F**) represents one cell or mouse.

**fig. S9. Tumor weight and immune cell infiltration in tumor microenvironment.** (**A** and **B**) Tumors from the experiment depicted in Fig. 5G (**A**) and Fig. 5J (**B**) were weighed and immune cell infiltration was assessed by flow cytometry. All data are means ± s.e.m. ns, not significant; \*\**p* < 0.01, \*\*\**p* < 0.001 and \*\*\*\**p* < 0.0001. Results in (**A** and **B**) are pooled from 2 independent experiments. Each symbol in (**A** and **B**) represents one mouse.

**fig. S10. Characterization of human SIRPα antibodies.** (**A** and **B**) Representative confocal microscopy images (**A**) and compiled data from 3 independent PLAs (**B**) with human SIRPα and human CD18, using human blood-derived macrophages, as was done for Fig. 3, J and K. Scale bar, 10 μm. (**C**) Compiled data from 3 independent binding assays with soluble human CD47-Fc and EL-4 cells expressing human SIRPα V1 or V2, in the presence of mAbs. (**D**) Compiled data from 3 independent binding assays with human SIRPα mAbs and the indicated fusion proteins, using ELISA. All data are means ± s.e.m. ns, not significant; \*\*\**p* < 0.001 and \*\*\*\**p* < 0.0001. Results in (**A**) are representative of 3 independent experiments. Results in (**B** to **D**) are pooled from 3 independent experiments. Each symbol in (**B**) represents one healthy donor.

**fig. S11. Impact of SIRPα-CD47 and SIRPα-CD18 interactions on phagocytosis and anti-tumor activity of macrophages.** SIRPα Abs disrupting the SIRPα-CD47 and SIRPα-CD18 interactions caused a greater increase in phagocytosis and better tumor suppression, compared to mAbs that primarily interrupted either interaction.

