## Supplementary Figures for "Binding of inhibitory checkpoints to CD18 in *cis* hinders anti-cancer immune responses"

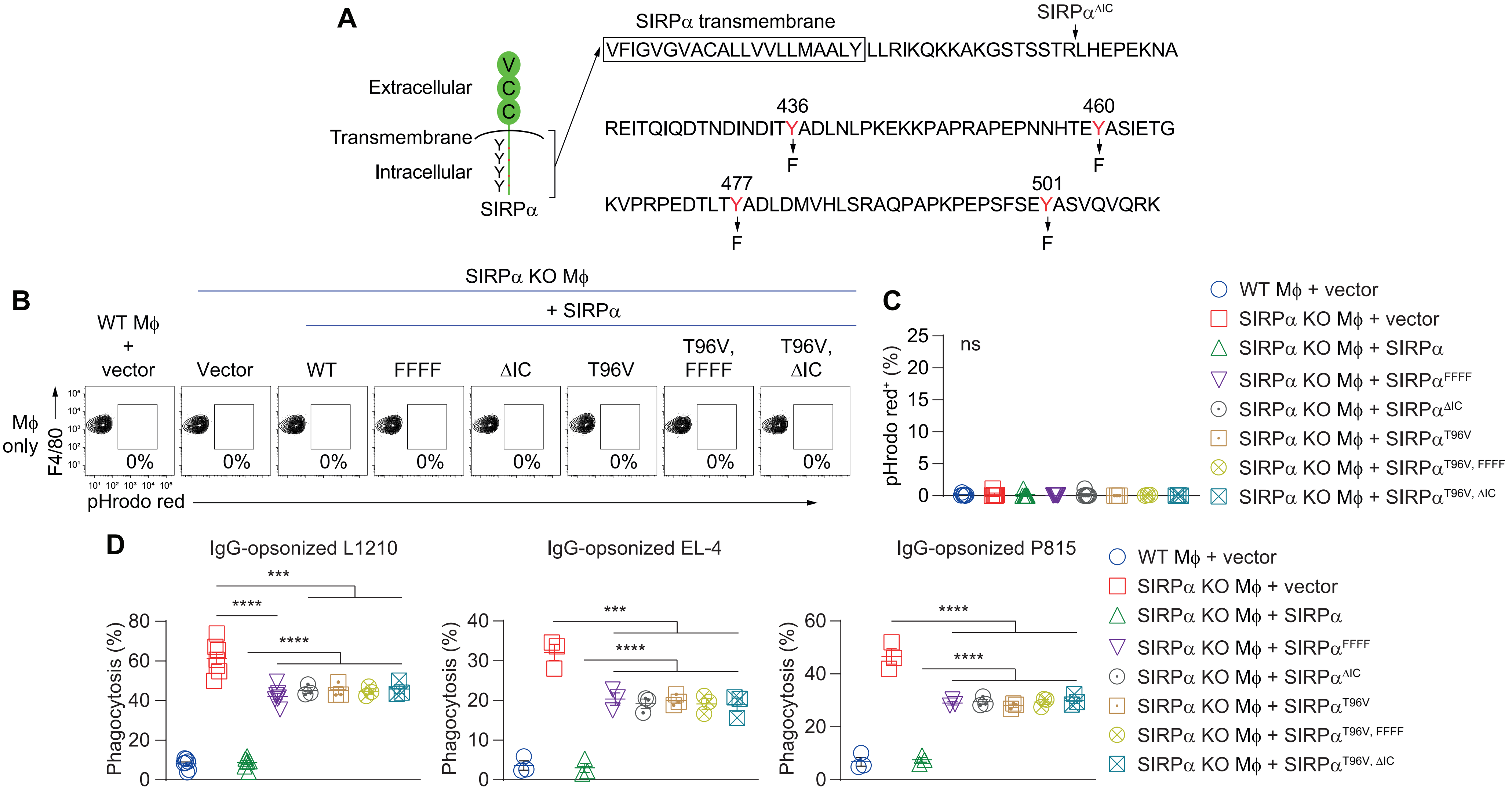
A
SIRPα transmembrane
SIRPα^ΔIC
VFIGVGVACALLVVLLMAALY LLRIKQKKAKGSTSSTRLHEPEKNA
Extracellular
Transmembrane
Intracellular
SIRPα
436
460
REITQIQDTNDINDITYADLNLPKEKKPAPRAPEPNNHTEYASIETG
F
F
477
501
KVPRPEDTLTYADLDMVHLSRAQPAPKPEPSFSEYASVQVQRK
F
F
B
SIRPα KO Mϕ
+ SIRPα
WT Mϕ + vector
Vector
WT
FFFF
ΔIC
T96V
T96V, FFFF
T96V, ΔIC
Mϕ only
F4/80
0%
pHrodo red
C
pHrodo red+ (%)
ns
WT Mϕ + vector
SIRPα KO Mϕ + vector
SIRPα KO Mϕ + SIRPα
SIRPα KO Mϕ + SIRPα^FFFF
SIRPα KO Mϕ + SIRPα^ΔIC
SIRPα KO Mϕ + SIRPα^T96V
SIRPα KO Mϕ + SIRPα^T96V, FFFF
SIRPα KO Mϕ + SIRPα^T96V, ΔIC
D
IgG-opsonized L1210
IgG-opsonized EL-4
IgG-opsonized P815
Phagocytosis (%)
***
****

Fig. S1

**A**

| Protein | Full name | Function | Normalized TIC | |
|---|---|---|---|---|
| | | | WT | SIRPα KO |
| SIRPα | Signal-regulatory protein α | Inhibitory receptor | 1.02E+07 | 0.00E+00 |
| CD45 | Protein tyrosine phosphatase, receptor type, C | Activating receptor | 7.08E+06 | 0.00E+00 |
| CD18 | Integrin β2 | Activating receptor | 5.38E+06 | 0.00E+00 |
| CD11b | Integrin αM | Activating receptor | 4.76E+06 | 0.00E+00 |
| ATP1A1 | Sodium/potassium-transporting ATPase subunit α 1 | sodium/potassium transporter | 8.25E+05 | 0.00E+00 |
| EMR1 | EGF-like module-containing mucin-like hormone receptor-like 1 | G-protein coupled receptor | 1.90E+05 | 0.00E+00 |
| JIP4 | C-jun-amino-terminal kinase-interacting protein 4 | Scaffold protein | 7.92E+06 | 0.00E+00 |
| AMPD2 | Adenosine monophosphate deaminase 2 | Enzyme | 1.46E+06 | 0.00E+00 |
| C1QC | Complement C1q C Chain | Complement protein | 1.31E+06 | 0.00E+00 |

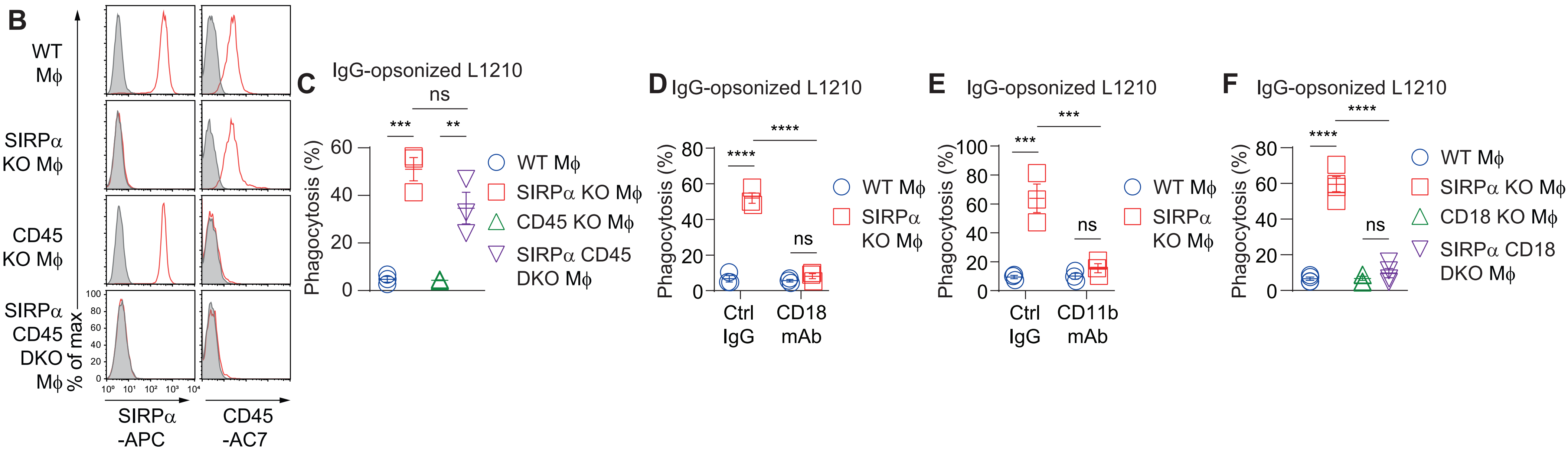


Fig. S2

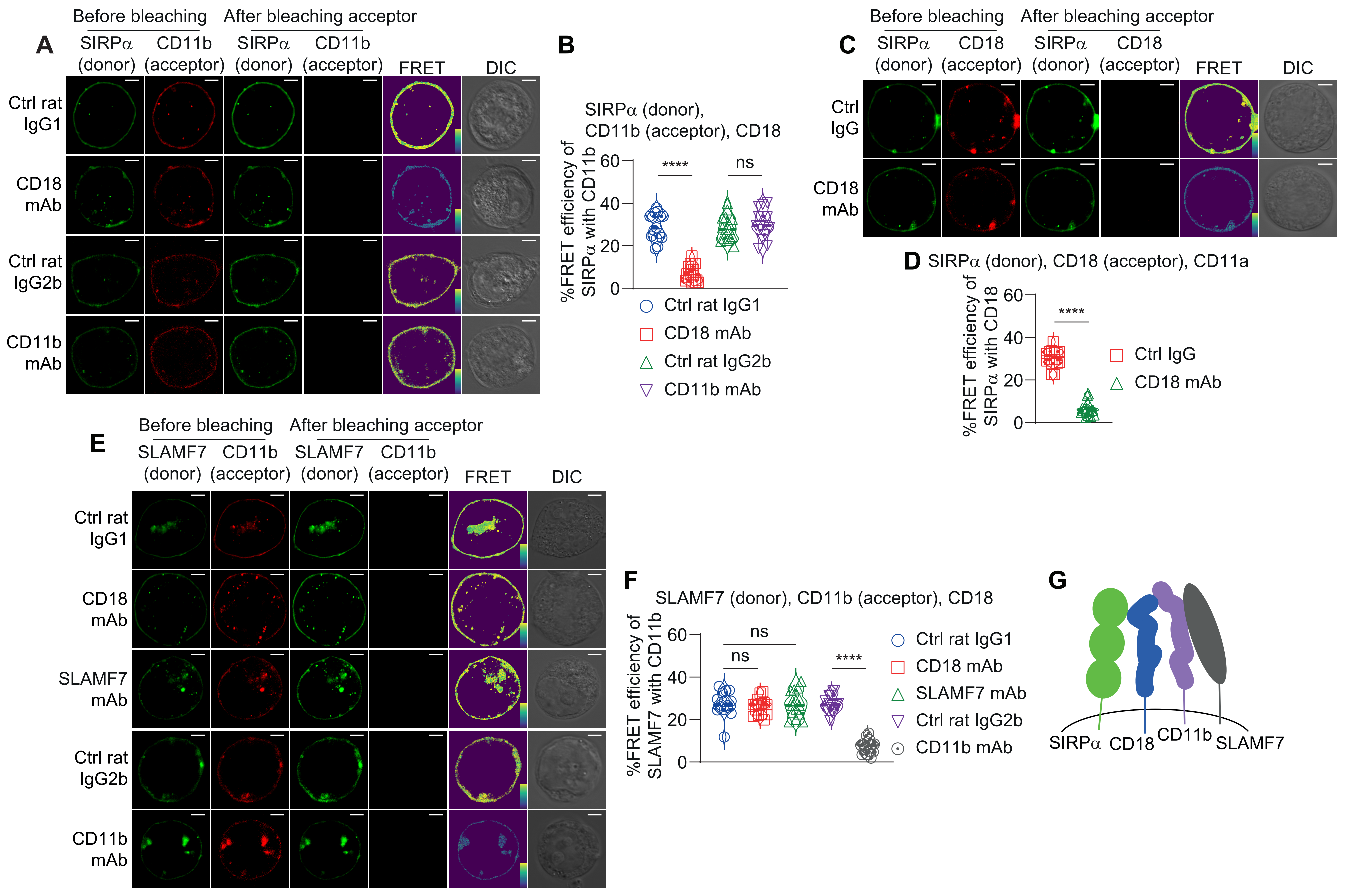
A
Before bleaching
After bleaching acceptor
SIRPα (donor)
CD11b (acceptor)
FRET
DIC
Ctrl rat IgG1
CD18 mAb
Ctrl rat IgG2b
CD11b mAb
B
SIRPα (donor), CD11b (acceptor), CD18
%FRET efficiency of SIRPα with CD11b
****
ns
Ctrl rat IgG1
CD18 mAb
Ctrl rat IgG2b
CD11b mAb
C
SIRPα (donor)
CD18 (acceptor)
Ctrl IgG
CD18 mAb
D
SIRPα (donor), CD18 (acceptor), CD11a
%FRET efficiency of SIRPα with CD18
Ctrl IgG
CD18 mAb
E
SLAMF7 (donor)
CD11b (acceptor)
SLAMF7 mAb
F
SLAMF7 (donor), CD11b (acceptor), CD18
%FRET efficiency of SLAMF7 with CD11b
G
SIRPα
CD18
CD11b
SLAMF7

Fig. S3

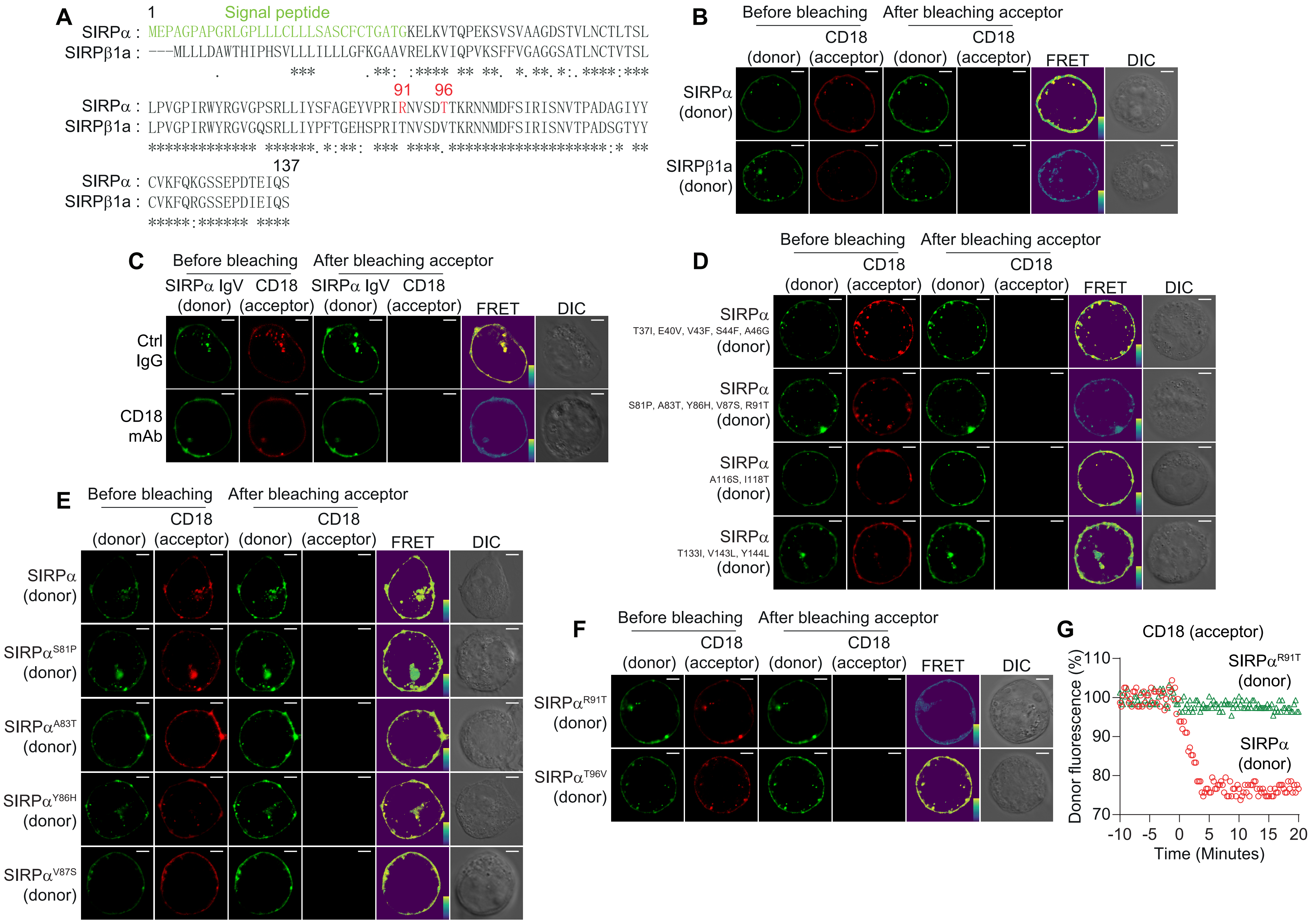
A
1
Signal peptide
SIRPα : MEPAGPAPGRLGPLLLCLLLSASCFCTGATGKELKVTQPEKSVSVAAGDSTVLNCTLTSL
SIRPβ1a : ---MLLLDAWTHIPHSVLLLILLLGFKGAAVRELKVIQPVKSFFVGAGGSATLNCTVTSL
91
96
SIRPα : LPVGPIRWYRGVGPSRLLIYSFAGEYVPRIRNVSDTTKRNNMDFSIRISNVTPADAGIYY
SIRPβ1a : LPVGPIRWYRGVGQSRLLIYPFTGEHSPRITNVSDVTKRNNMDFSIRISNVTPADSGTYY
137
SIRPα : CVKFQKGSSEPDTEIQS
SIRPβ1a : CVKFQRGSSEPDIEIQS
B
Before bleaching
After bleaching acceptor
CD18
(donor)
(acceptor)
FRET
DIC
SIRPα (donor)
SIRPβ1a (donor)
C
SIRPα IgV
Ctrl IgG
CD18 mAb
D
SIRPα T37I, E40V, V43F, S44F, A46G (donor)
SIRPα S81P, A83T, Y86H, V87S, R91T (donor)
SIRPα A116S, I118T (donor)
SIRPα T133I, V143L, Y144L (donor)
E
SIRPα (donor)
SIRPα^S81P (donor)
SIRPα^A83T (donor)
SIRPα^Y86H (donor)
SIRPα^V87S (donor)
F
SIRPα^R91T (donor)
SIRPα^T96V (donor)
G
CD18 (acceptor)
Donor fluorescence (%)
Time (Minutes)
SIRPα^R91T (donor)
SIRPα (donor)
110
100
90
80
70
-10
-5
0
5
10
15
20

Fig. S4

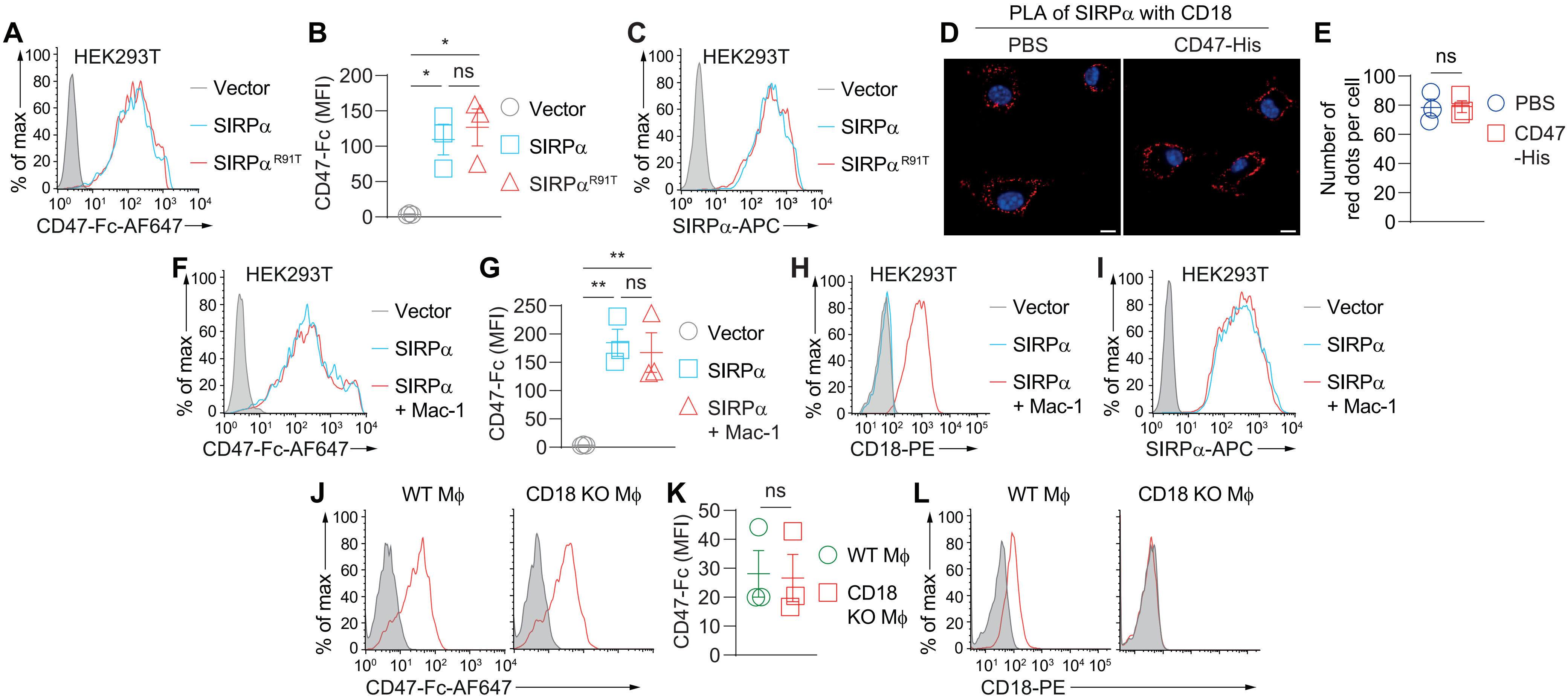

A
HEK293T
% of max
CD47-Fc-AF647
Vector
SIRPα
SIRPα R91T
B
CD47-Fc (MFI)
*
ns
C
SIRPα-APC
D
PLA of SIRPα with CD18
PBS
CD47-His
E
Number of red dots per cell
CD47 -His
F
SIRPα + Mac-1
G
**
H
CD18-PE
I
J
WT Mϕ
CD18 KO Mϕ
K
L


Fig. S5

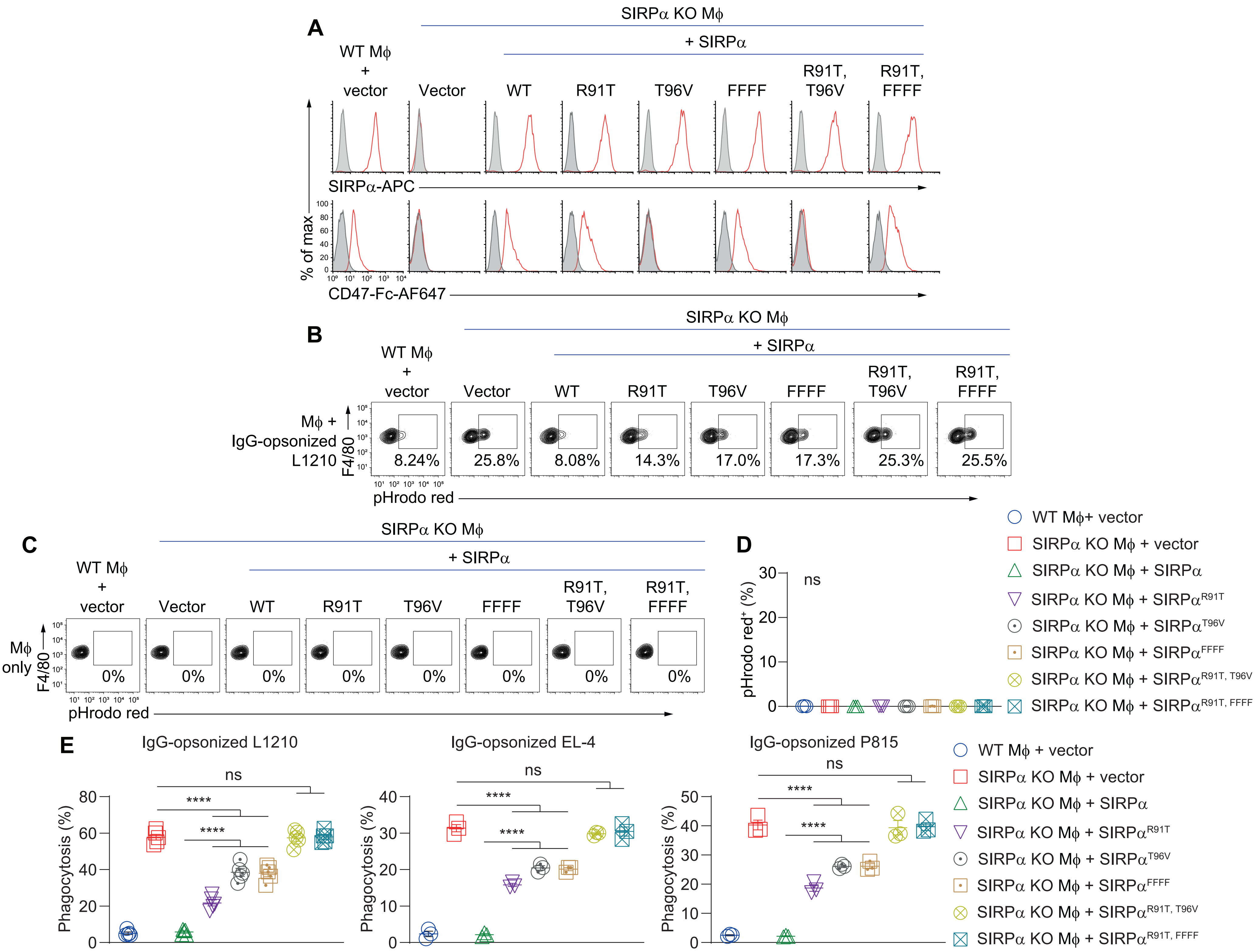

A
SIRPα KO Mϕ
+ SIRPα
WT Mϕ + vector
Vector
WT
R91T
T96V
FFFF
R91T, T96V
R91T, FFFF
SIRPα-APC
% of max
CD47-Fc-AF647
B
SIRPα KO Mϕ
+ SIRPα
WT Mϕ + vector
Vector
WT
R91T
T96V
FFFF
R91T, T96V
R91T, FFFF
Mϕ + IgG-opsonized L1210
F4/80
8.24%
25.8%
8.08%
14.3%
17.0%
17.3%
25.3%
25.5%
pHrodo red
C
SIRPα KO Mϕ
+ SIRPα
WT Mϕ + vector
Vector
WT
R91T
T96V
FFFF
R91T, T96V
R91T, FFFF
Mϕ only
F4/80
0%
0%
0%
0%
0%
0%
0%
0%
pHrodo red
D
pHrodo red+ (%)
ns
WT Mϕ+ vector
SIRPα KO Mϕ + vector
SIRPα KO Mϕ + SIRPα
SIRPα KO Mϕ + SIRPαR91T
SIRPα KO Mϕ + SIRPαT96V
SIRPα KO Mϕ + SIRPαFFFF
SIRPα KO Mϕ + SIRPαR91T, T96V
SIRPα KO Mϕ + SIRPαR91T, FFFF
E
IgG-opsonized L1210
IgG-opsonized EL-4
IgG-opsonized P815
Phagocytosis (%)
ns
****
WT Mϕ + vector
SIRPα KO Mϕ + vector
SIRPα KO Mϕ + SIRPα
SIRPα KO Mϕ + SIRPαR91T
SIRPα KO Mϕ + SIRPαT96V
SIRPα KO Mϕ + SIRPαFFFF
SIRPα KO Mϕ + SIRPαR91T, T96V
SIRPα KO Mϕ + SIRPαR91T, FFFF


Fig. S6

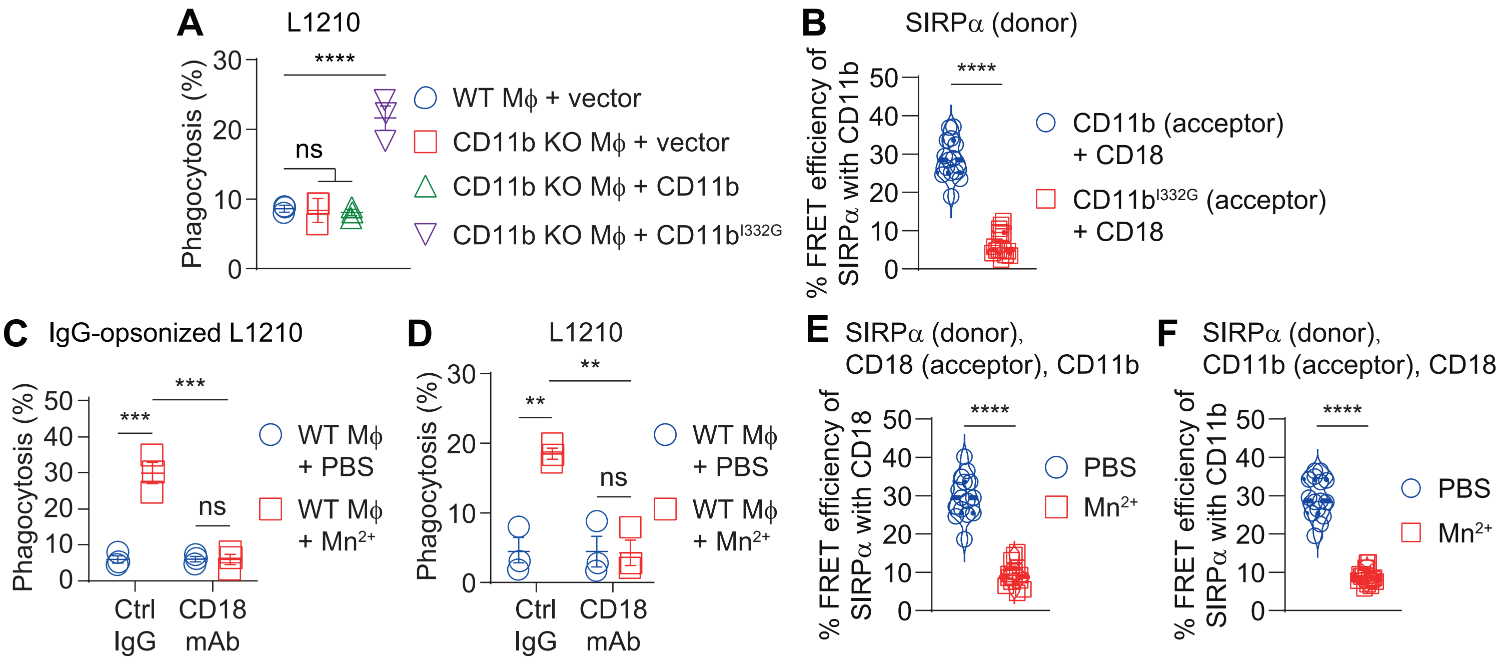
A
L1210
Phagocytosis (%)
****
ns
WT Mϕ + vector
CD11b KO Mϕ + vector
CD11b KO Mϕ + CD11b
CD11b KO Mϕ + CD11b^I332G
B
SIRPα (donor)
% FRET efficiency of SIRPα with CD11b
****
CD11b (acceptor) + CD18
CD11b^I332G (acceptor) + CD18
C
IgG-opsonized L1210
Phagocytosis (%)
***
***
ns
Ctrl IgG
CD18 mAb
WT Mϕ + PBS
WT Mϕ + Mn2+
D
L1210
Phagocytosis (%)
**
**
ns
Ctrl IgG
CD18 mAb
WT Mϕ + PBS
WT Mϕ + Mn2+
E
SIRPα (donor), CD18 (acceptor), CD11b
% FRET efficiency of SIRPα with CD18
****
PBS
Mn2+
F
SIRPα (donor), CD11b (acceptor), CD18
% FRET efficiency of SIRPα with CD11b
****
PBS
Mn2+

Fig. S7

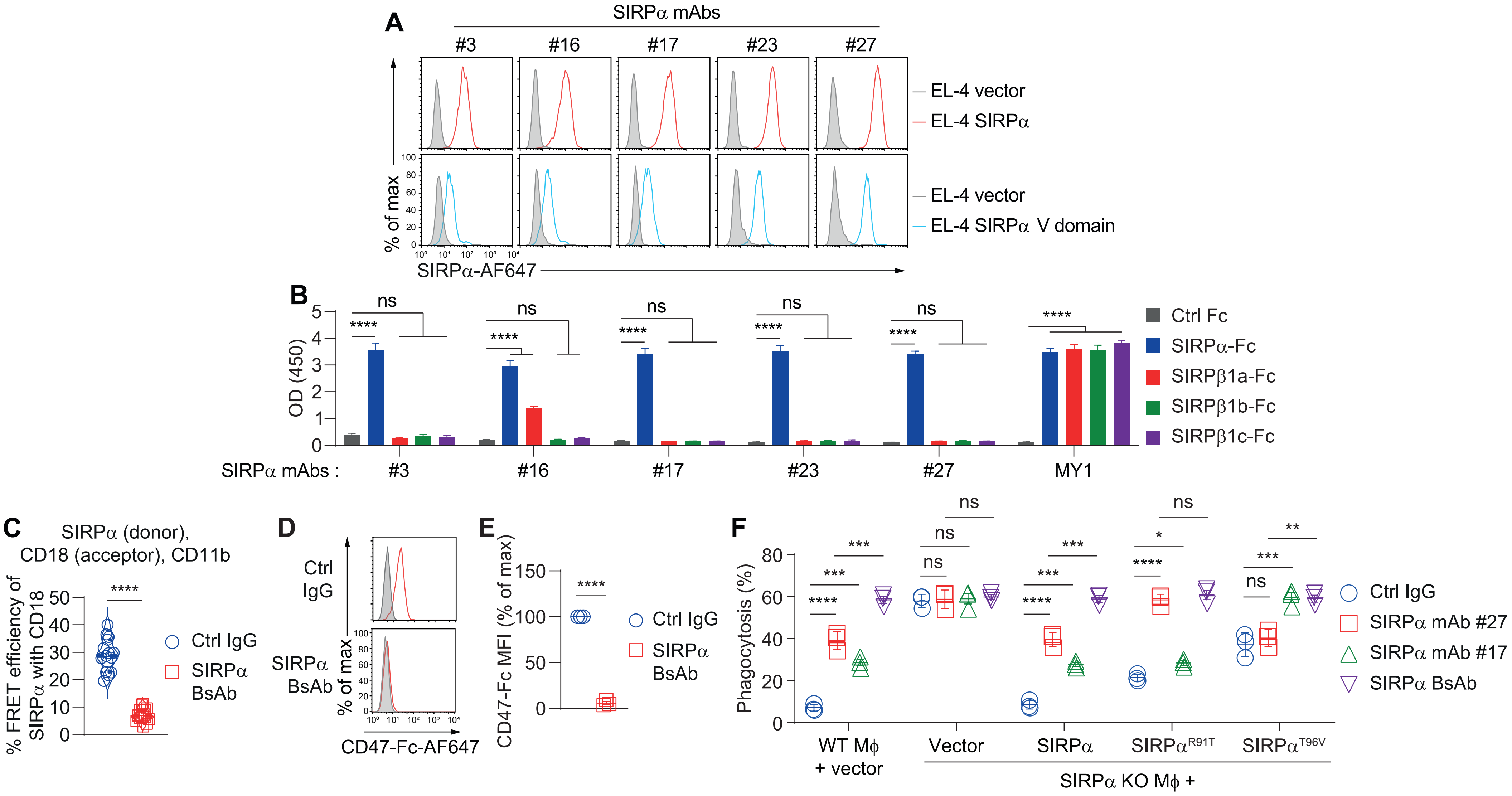
A
SIRPα mAbs
#3
#16
#17
#23
#27
EL-4 vector
EL-4 SIRPα
EL-4 vector
EL-4 SIRPα V domain
% of max
SIRPα-AF647
B
OD (450)
SIRPα mAbs :
MY1
Ctrl Fc
SIRPα-Fc
SIRPβ1a-Fc
SIRPβ1b-Fc
SIRPβ1c-Fc
ns
****
C
SIRPα (donor),
CD18 (acceptor), CD11b
% FRET efficiency of SIRPα with CD18
Ctrl IgG
SIRPα BsAb
D
Ctrl IgG
SIRPα BsAb
% of max
CD47-Fc-AF647
E
CD47-Fc MFI (% of max)
Ctrl IgG
SIRPα BsAb
F
Phagocytosis (%)
WT Mϕ + vector
Vector
SIRPα
SIRPα R91T
SIRPα T96V
SIRPα KO Mϕ +
Ctrl IgG
SIRPα mAb #27
SIRPα mAb #17
SIRPα BsAb

Fig. S8

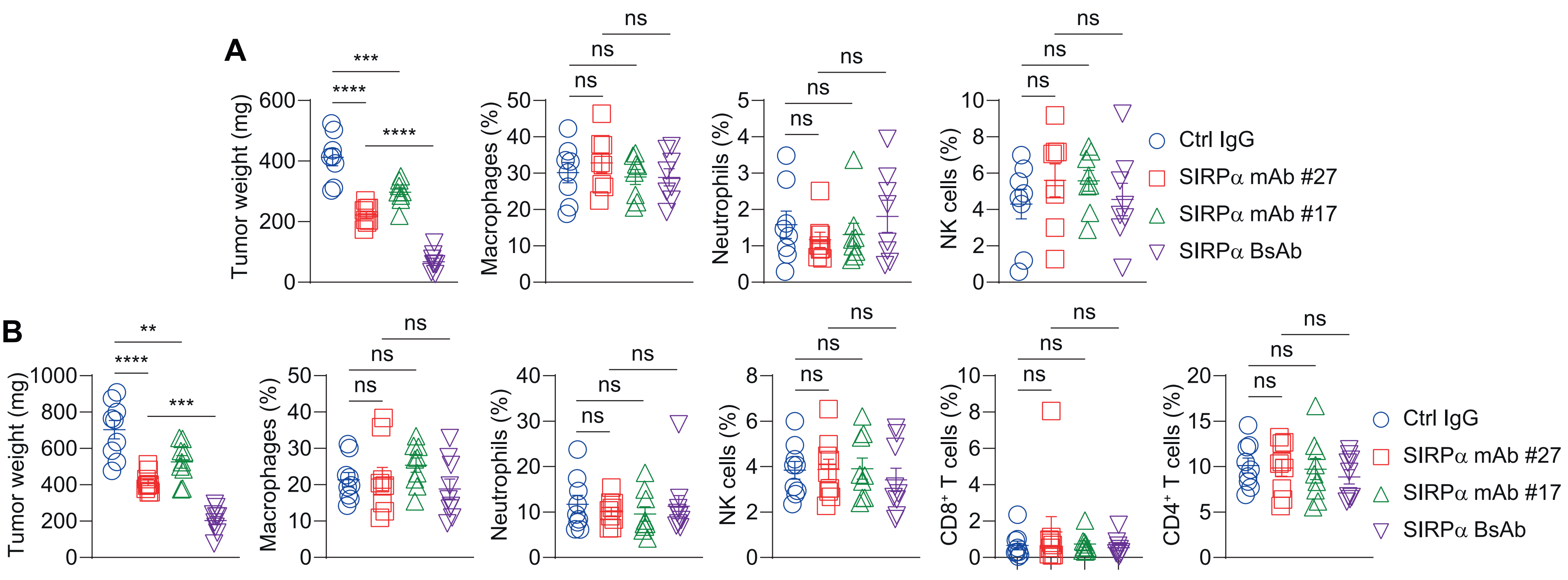
A
B
Tumor weight (mg)
Macrophages (%)
Neutrophils (%)
NK cells (%)
CD8+ T cells (%)
CD4+ T cells (%)
ns
***
****
**
Ctrl IgG
SIRPα mAb #27
SIRPα mAb #17
SIRPα BsAb

Fig. S9

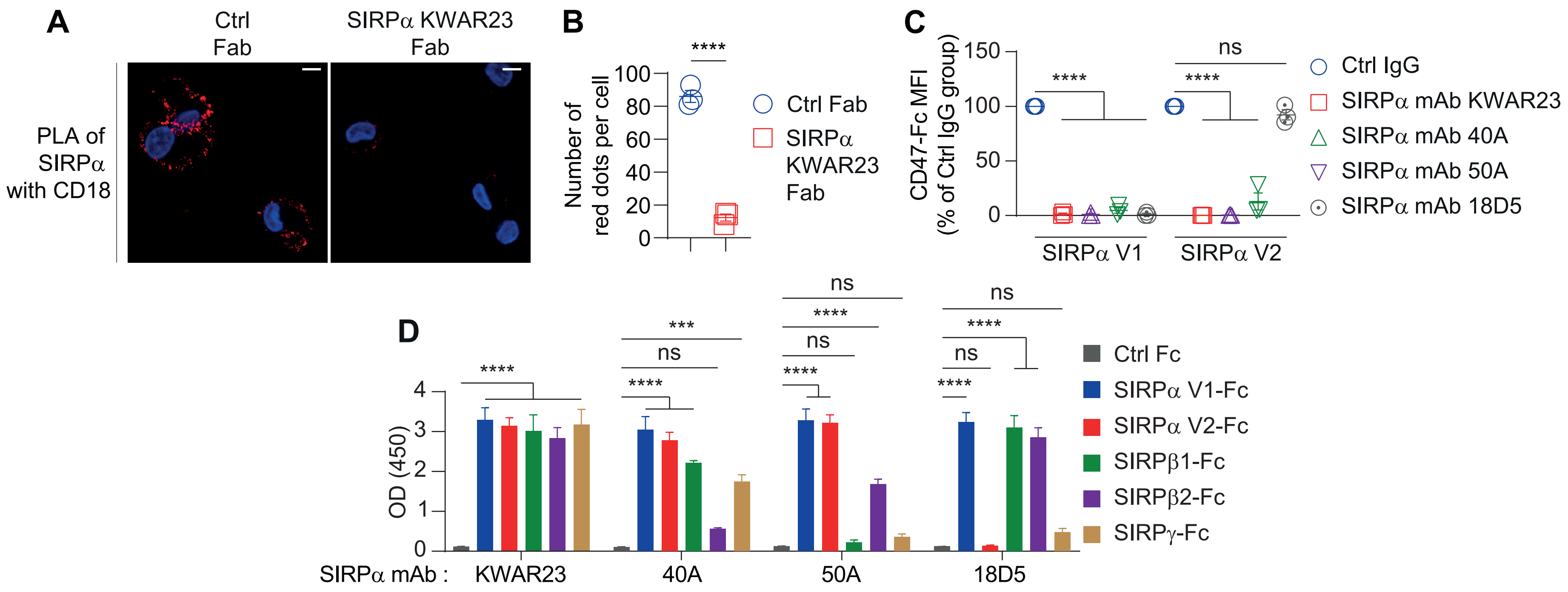
A
Ctrl Fab
SIRPα KWAR23 Fab
PLA of SIRPα with CD18
B
Number of red dots per cell
****
Ctrl Fab
SIRPα KWAR23 Fab
C
CD47-Fc MFI (% of Ctrl IgG group)
****
ns
SIRPα V1
SIRPα V2
Ctrl IgG
SIRPα mAb KWAR23
SIRPα mAb 40A
SIRPα mAb 50A
SIRPα mAb 18D5
D
OD (450)
SIRPα mAb :
KWAR23
40A
50A
18D5
Ctrl Fc
SIRPα V1-Fc
SIRPα V2-Fc
SIRPβ1-Fc
SIRPβ2-Fc
SIRPγ-Fc

Fig. S10

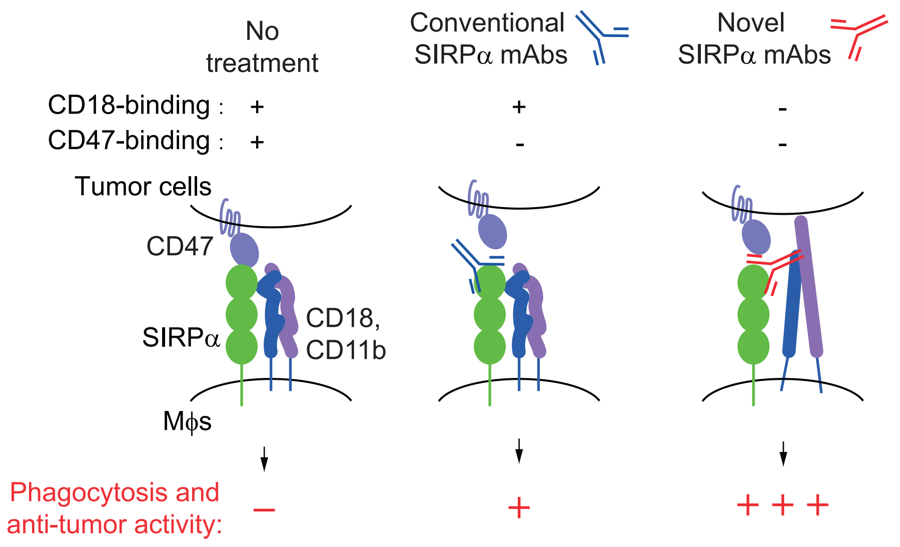
No treatment
Conventional SIRPα mAbs
Novel SIRPα mAbs
CD18-binding : + + -
CD47-binding : + - -
Tumor cells
CD47
SIRPα
CD18, CD11b
Mϕs
Phagocytosis and anti-tumor activity: – + + + +

Fig. S11
